# DNA methylation reveals molecular heterogeneity and urate excretion-associated subtypes of gout

**DOI:** 10.64898/2026.10.19.760079

**Authors:** Guangmin Zheng, Aichang Ji, Wenrui Shi, Xinde Li, Xiuyan Ruan, Xiaopeng Ji, Xiangdong Fang, Changgui Li, Hongzhu Qu

**Author notes:** **Corresponding author:** Hongzhu Qu, Changgui Li, Xiangdong Fang. These authors contributed equally (Guangmin Zheng, Aichang Ji).

## Abstract

**Background:** The molecular heterogeneity among gout patients remains largely unexplored, hindered the precise treatment for gout. Common indices for gout subtyping are cumbersome to measure, resulting in poor patient compliance.

**Methods:** We elucidated the molecular heterogeneity of gout and developed a simple, reliable classification method for gout subtypes based on genome-wide DNA methylation, to facilitate individualized treatment. Genome-wide DNA methylation profiling was conducted in two independent cohorts, comprising a discovery cohort (n = 108) and a validation cohort (n = 100) representing four clinical gout subtypes.

**Results:** We identified two gout subgroups associated with the fractional excretion of uric acid (FE_UA_) but not with 24-hour urinary uric acid levels (UUE) based on unsupervised clustering of whole-genome DNA methylation profiles, highlighting the contribution of epigenetic regulation to molecular heterogeneity in urate metabolism. Enrichment analysis links FE_UA_-associated subtypes to renal cell types, calcium related pathways and extended to the circulating immune compartment, with neutrophil- versus memory B cell-skewed leukocyte composition. A methylation-based classification model using 18 LASSO-selected CpGs with an XGBoost classifier effectively distinguished patients with high and low uric acid excretion, achieving excellent predictive performance (training cohort: AUC = 0.96, AP = 0.91; validation cohort: AUC = 0.86, AP = 0.85), outperforming the published model with clinical characteristics and SNPs. The signature site-associated genes *NEK1* and *TRPS1* display significant expression differences among urate-stratified patients and immune cells groups. *TRPS1* may affect the AP-1/NF-κB inflammatory programme.

**Conclusions:** These findings reveal distinct DNA methylation signatures underlying gout heterogeneity and demonstrate the feasibility of DNA methylation markers for gout subtype classification. The proposed methylation-based model provides a promising approach for personalized management of gout.

## Background

Gout is a chronic metabolic inflammatory arthritis driven by abnormal uric acid (UA) metabolism and monosodium urate (MSU) deposition, leading to severe joint and renal complication[1]. Therefore, rigorous monitoring of serum urate levels and effective urate-lowering therapy (ULT) are central to gout management, aiming for serum urate levels below 6 mg/dL[2, 3]. Although current guidelines do not recommend ULT selection based on hyperuricemia subtype due to limited evidence, increasing data support individualized management based on four hyperuricemia subtypes: [4–6]: renal urate overload (ROL), underexcretion (RUE), combined (RCB), and normal (RUN)[7, 8]. Accurately distinguishing urate overproduction from underexcretion before initiating ULT is critical, as uricosuric agents can exacerbate crystal formation and renal damage in overproducers [9]. Thus, an accurate assessment of urate excretion patterns prior to initiating ULT is vital for safe and effective gout management.

Accurate gout subtyping relies on the precise assessment of FE_UA_ and UUE. The established gold standard requires a 24-hour urine collection, preceded by a 14-day medication washout and a 5-day purine-restricted diet[10]. Because many drugs influence renal urate excretion, discontinuing all medications is required but often clinically precarious. Moreover, the inherent inconvenience of 24-hour urine collection and strict dietary control makes this procedure operationally complex and time-consuming. This greatly limits its practicality in routine practice and prone to poor compliance, especially in children and critically ill individuals[11]. Although spot urine or short-term samples have been explored as alternatives[12–14], practical barriers remain. Consequently, a simple and reliable subtyping method is needed to support clinical decision-making.

Epigenetic regulation links between genetic predisposition and environmental exposures in complex diseases such as gout[15]. Among these mechanisms, DNA methylation provides a stable, quantifiable link between environmental factors and disease phenotypes[16]. While methylation-based classifiers have shown great success in oncology, such as accurate molecular subtyping and improved prognostic prediction [17, 18], systematic studies defining gout molecular subtypes remain limited, despite evidences of distinct methylation patterns between gout patients and healthy controls were reported[19–21]. Since direct sampling of kidney and intestine, which are the primary sites of UA metabolism, in gout patients is rarely clinically warranted, peripheral blood mononuclear cells (PBMC) provide an accessible alternative to capture systemic molecular alterations[22]. Consequently, genome-wide DNA methylation profiling of PBMCs serves as a practical approach to investigate epigenetic changes in gout.

To address this gap, whole-genome bisulfite sequencing (WGBS) was conducted on PBMC samples to profile the DNA methylome. By integrating unsupervised clustering, differential methylation, and machine learning, we identified subtype-specific methylation signatures underlying gout heterogeneity. We subsequently developed a methylation-based classifier to accurately predict urate excretion subtypes, offering a simplified and standardized molecular framework to advance precision management in gout.

## Methods

### Cohort description and sample collection

We enrolled male patients diagnosed with gout according to the 2015 ACR/EULAR classification criteria[23]. Exclusion criteria included severe comorbidities, severe metabolic abnormalities (blood pressure ≥180/110 mmHg, fasting blood glucose ≥ 11.1 mmol/L), renal impairment (eGFR <45 mL/min/1.73 m²), and regular anticoagulant use. The study comprised a discovery cohort (n=108; recruited July 2016–March 2019) and a validation cohort (n=100; recruited December 2019–June 2023) from the Affiliated Hospital of Qingdao University. Clinical characteristics and subtype distributions are summarized in **Supplementary Tables 1 and 2**, respectively. The study protocol was approved by the local Ethics Committee (QYFYKYLL923011921) and complied with the Declaration of Helsinki. All participants provided written informed consent.

### Clinical assessment and gout subtyping

Following a 14-day medication washout and a 5-day low-purine diet (<200 mg/day), 24-hour urine and fasting whole blood samples (stored at −80 °C) were collected[10]. Clinical data were extracted from the hospital electronic health record. Serum uric acid (sUA), blood glucose, triglycerides, total cholesterol, low-density lipoprotein, high-density lipoprotein, blood urea nitrogen, serum creatinine (sCr), urinary uric acid (uUA), and urine creatinine (uCr) were measured using an automated biochemical analyzer (TBA-40FR; Toshiba, Japan). Fractional excretion of uric acid (FE_UA_) was calculated as (FE_UA_ = uUA/uCr × sCr/sUA). Gout subtypes—renal urate underexcretion (RUE), overload (ROL), combined (RCB), and normal (RUN)—were classified using cutoffs of 600 mg/d/1.73 m² for urinary uric acid excretion (UUE) and 5.5% for FE_UA_.

### Genotyping and WGBS

In the model comparison, the machine learning model for underexcretion-type gout, incorporating six clinical features and four replicated genetic loci, was developed based on our previous work[24]. The discovery cohort in this study is a subset dataset of this previously publication, and details regarding the genotyping procedures can be found therein. For DNA methylation, genomic DNA (gDNA) was extracted from PBMCs using a magnetic bead-based method according to the manufacturer’s instructions. 500μg of purified genomic DNA underwent bisulfite conversion using the EZ DNA Methylation-Gold™ Kit. WGBS libraries were constructed and sequenced on the Illumina NovaSeq 6000 platform at a targeted 30× depth with a ≥99% cytosine-to-thymine conversion rate.

### Statistical analysis

Continuous variables are presented as medians with interquartile ranges, and categorical variables as frequencies. Group comparisons were evaluated using the *Wilcoxon* test for continuous data and *Fisher*’s exact test for categorical data. A two-sided *P*-value < 0.05 was considered statistically significant.

### Cell-type-specific virtual knockout analysis of *NEK1* and *TRPS1*

Cell-type-specific virtual knockout (vKO) analysis was performed using scTenifoldKnk v1.1.5 / scTenifoldNet v1.4.3 (R 4.5.3) [25] based on the scRNA-seq data. Healthy-control cells were used as the unperturbed wild-type reference. Based on their predominant expression patterns, *NEK1* was virtually knocked out in CD8□ T, whereas *TRPS1* was knocked out in monocyte–macrophages. Cell-type-specific gene regulatory networks were constructed, and pseudo-knockout networks were generated by removing the outgoing regulatory edges of the target gene. Differentially regulated genes between the wild-type and pseudo-knockout networks were identified through manifold alignment and used for downstream functional enrichment analysis. Cell subsampling and tensor initialization are stochastic, so the pipeline was repeated with seed = 1, 2, 3 and 42. For *TRPS1* vKO, the 62 genes significant in all four runs (72 %) were carried forward; 47 were ribosomal/translation genes, leaving 14 besides *TRPS1*. In contrast, *NEK1* vKO did not yield a reproducible downstream module in CD8□ T cells, likely because its low detection rate prevented reliable inference of its outgoing regulatory edges. As scTenifoldKnk simulates knockout by removing these inferred edges, the *NEK1* result was considered inconclusive and was not included in downstream enrichment analysis.

### STRING network and gene set enrichment

For *TRPS1* vKO, the 14 genes were queried against STRING v12.0 at combined score thresholds 0.4 and 0.15; scores ≥ 0.4 are referred to as high-to-medium confidence and 0.15–0.4 as sub-threshold. Genes were ranked by mean Z score across seeds. Because the ranking was dominated by ribosomal-protein genes, ribosomal/translation genes were removed. GSEA used fgsea v1.36.2 against 5,067 sets from MSigDB H, C5 GO:BP, C2 CP:REACTOME and C2 CP:KEGG_LEGACY (msigdbr v26.1.1), restricted to tested genes.

### Supplementary Materials

Additional methodological details are available in the Supplementary Material. Figs. S1 to S5. Tables S1 to S10.

## Results

### Genes linked to DMSs in RUN gout patients are enriched in the regulation of membrane potential

We first examined DNA methylation patterns across the four clinical subtypes in the discovery cohort (n = 108) and observed no prominent group-level differences. When identifying the differentially methylated loci, samples were treated as the case group for each subtype while the remaining three subtypes served as the control group. The variations in differential methylation features among the four subtypes were minimal (**Fig. 1A**). Moreover, principal component analysis (PCA) based on the differentially methylation sites (DMSs) failed to separate the subtypes distinctly (**Fig. 1B**). Patients classified as RCB and RUE appeared closer to each other, while ROL and RUN patients clustered together. These results suggest a clear separability between the RCB/RUE and ROL/RUN groups in terms of differential methylation features, highlighting a discrepancy between clinically defined subtypes and intrinsic molecular patterns. Given this, we hypothesized that a four-class DNA methylation-based classifier would exhibit limited predictive performance. To evaluate this, we constructed a multiclass model using the differential methylation features from all four groups. Among the 108 patients in the discovery cohort, samples were randomly divided into training and testing sets at a 7:3 ratio, and a five-fold cross-validation strategy was applied. As shown in **Fig. S1A**, model performance on the testing set varied substantially across subtypes. Both the area under the curve (AUC) and average precision (AP) exhibited considerable variability, indicating instability in classification performance. Specifically, the weighted average AUC ranged from 0.80 to 0.93, while the weighted average AP ranged from 0.60 to 0.79. These results support our hypothesis that the discriminative capacity of a four-class model is limited.

**Figure 1.**
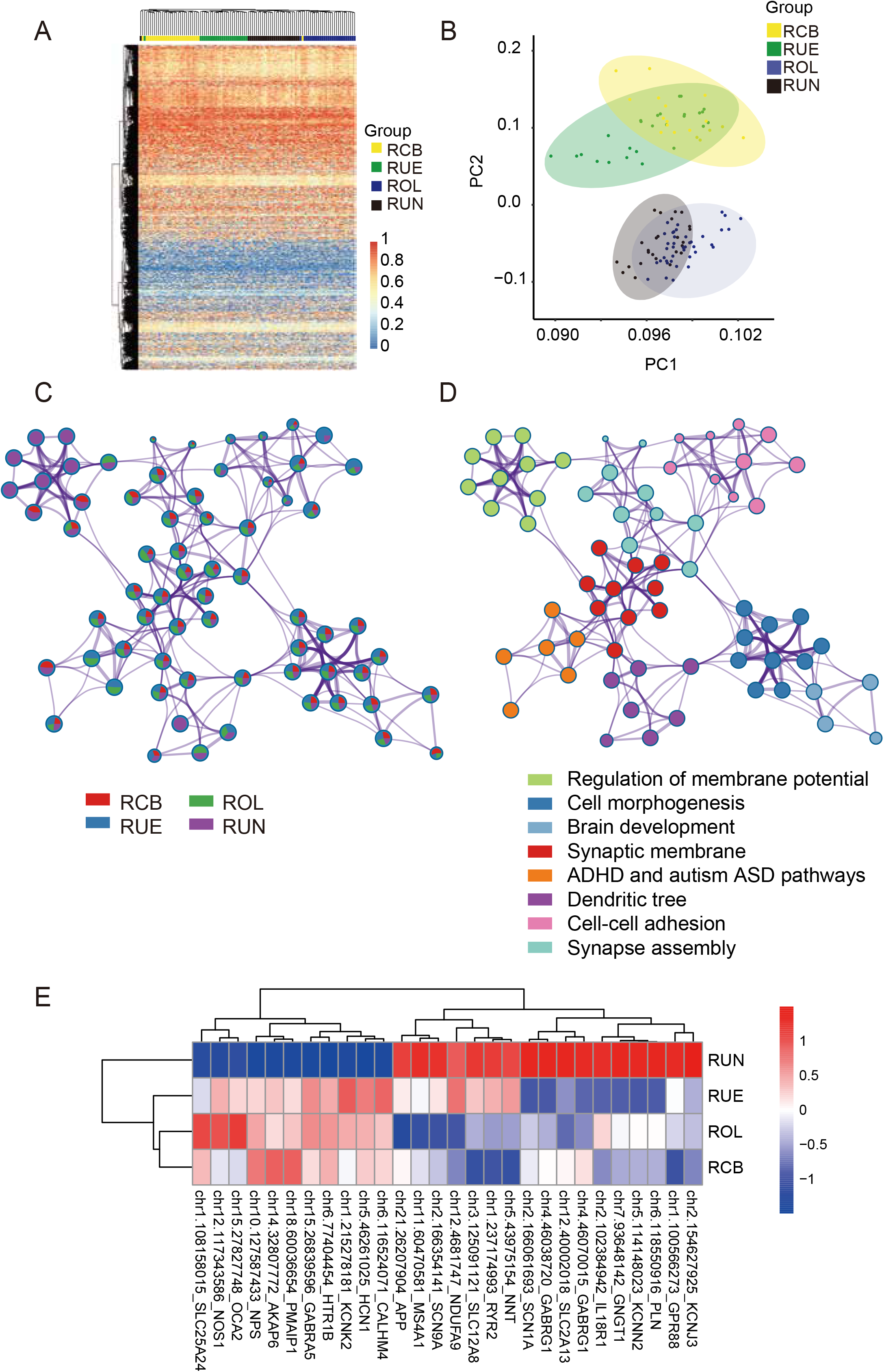
Differential methylation characteristics of four clinical gout subtypes. (A) Heatmap of differential methylation characteristics for the four clinical gout subtypes. For each subtype, samples were treated as the case group while the remaining subtypes served as the control group, and all resulting differential DNA methylation features were used for cross-group comparison. (B) PCA plot of differential DNA methylation features across the four clinical gout subtypes. (C) PPI network of genes annotated from DMSs, with different colors representing distinct gout subtypes. (D) PPI network of genes annotated from DMSs, with different colors indicating distinct protein functional modules. (E) DMSs and their associated genes are specifically enriched in the membrane potential regulation pathway in RUN gout patients. PCA, principal component analysis; PPI, protein-protein interaction; RCB, renal urate combined; RUE, renal urate underexcretion; ROL, renal urate overload; RUN, renal urate normal; DMSs, differentially methylated sites.

We further explored the differential methylation features and the specific protein interaction pathways regulated in the different clinical subtypes of gout. The analysis revealed substantial overlap of differentially methylated genes and enriched pathways across the four subtypes (**Fig. S1B, C**). The protein-protein interaction (PPI) networks of genes associated with these methylation sites were enriched in numerous common functional pathways, including cell morphology, brain development, synaptic membranes, attention-deficit/hyperactivity disorder, autism spectrum disorder, dendritic structures, cell adhesion, and synapse assembly across four subtypes (**Fig. 1C, D**), indicating a certain level of shared methylation-regulated mechanisms among the subtypes.

Interestingly, when investigating subtype-specific regulatory pathways, we found that the RUN type exhibited significant and specific enrichment of DMSs in the regulation of membrane potential pathway (purple nodes in **Fig. 1C** and green nodes in **Fig. 1D)**. These specific methylation sites showed a distinct pattern in the RUN subtype (**Fig. 1E**). This unique enrichment pattern suggests that the RUN subtype harbors distinct methylation features associated with membrane potential regulation, potentially reflecting its specific pathophysiological mechanisms. Such findings provide new insights into the molecular underpinnings of gout in patients with the RUN subtype.

### Unsupervised clustering of DNA methylation identifies FE_UA_-associated molecular subtypes of gout

Based on the above findings, we further investigated the molecular subtypes of gout patients independent of their clinical classifications, aiming to uncover intrinsic molecular drivers underlying gout heterogeneity. We first performed correlation analysis on genome-wide DNA methylation profiles from 108 gout patients and observed a high degree of inter-patient correlation, with most correlation coefficients exceeding 0.7 (**Fig. S2A**), indicating overall homogeneity and strong similarity in DNA methylation patterns among patients. Next, we performed unsupervised clustering on genome-wide DNA methylation profiles from these samples and detected two distinct clusters, designated Cluster1 and Cluster2 (**Fig. 2A**). This clustering pattern was consistently validated in the independent validation cohort of 100 gout patients, in which the optimal cluster number was also determined to be two (**Fig. S2B–C**).

**Figure 2.**
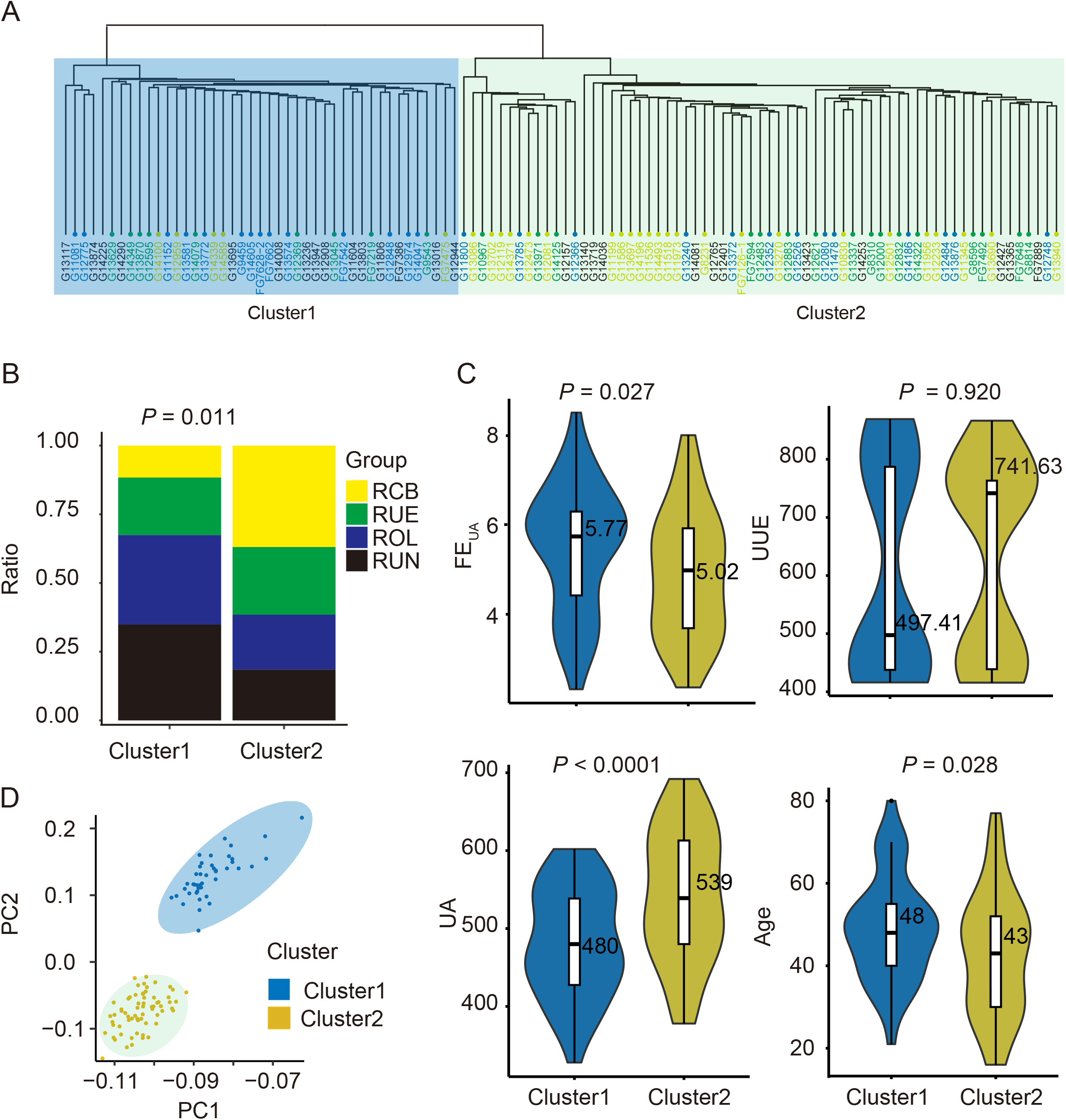
Unsupervised clustering analysis of gout patients in the discovery cohort. (A) Unsupervised clustering analysis of DNA methylation features. The blue and green background box stands for Cluster1 and Cluster2, respectively. (B) Correlation analysis between unsupervised clustering and clinical subtypes of gout, evaluated using *Fisher’s* exact test. (C) Comparative analysis of FE_UA_, UA, UUE, and age between the two clusters identified by unsupervised clustering. *P* values were calculated using the *Wilcoxon* rank sum test. (D) PCA of the top 2000 DMSs. FE_UA_, the fractional excretion of uric acid; UA, uric acid; UUE, 24-hour urinary urate excretion; PCA, principal component analysis. DMSs, differentially methylated sites.

To explore the relationship between the unsupervised clusters and clinical gout subtypes, we integrated clinical subtype information with clustering results for correlation and comparative analyses. The Cluster1 was associated with the ROL and RUN clinical subtypes, characterized by higher FE_UA_ and lower UA levels (**Fig. 2B-C**). Conversely, the Cluster2 correlated with the RCB and RUE subtypes, showing lower FE_UA_ and elevated UA levels. However, no significant difference was observed in UUE between the two clusters (**Fig. 2B-C**). These findings were consistently validated in the independent cohort (**Fig. S2D–E**). Further comparisons of other clinical parameters revealed a significant difference in age between the two clusters, whereas estimated glomerular filtration rate, serum creatinine, fasting glucose, and other variables showed no statistically significant differences (*P* > 0.05). In addition, PCA of the top 2000 most variable DMSs (**Fig. 2D**) demonstrated clear separation between the Cluster1 and Cluster2 clusters, further supporting the presence of distinct DNA methylation signatures between these molecular subtypes.

### Enrichment analysis links FE_UA_-associated subtypes to renal cell types and calcium related pathways

To further characterize the molecular differences between the two unsupervised clusters, we performed differential DNA methylation analysis and identified pronounced methylation differences between Cluster1 and Cluster2 (*q* value < 0.05 and |meth.diff| > 10%). Compared with Clusrer1, the Cluster2 exhibited higher methylation levels at hypermethylated sites and lower methylation levels at hypomethylated sites, with the overall differences largely driven by hypermethylation **(Fig. 3A** and **Fig. S3A)**. Cell-type enrichment analysis revealed that genes annotated to hypermethylated sites were significantly enriched in renal functional cell types, including vascular smooth muscle cells, pericytes, connecting tubule cells, collecting duct cells, and proximal tubular epithelial cells, as well as midbrain neurons (**Fig. 3B**). These findings suggest that peripheral blood DNA methylation alterations may reflect renal functional changes, consistent with our earlier findings that the strong associations between cluster, FE_UA_, and serum urate levels.

**Figure 3.**
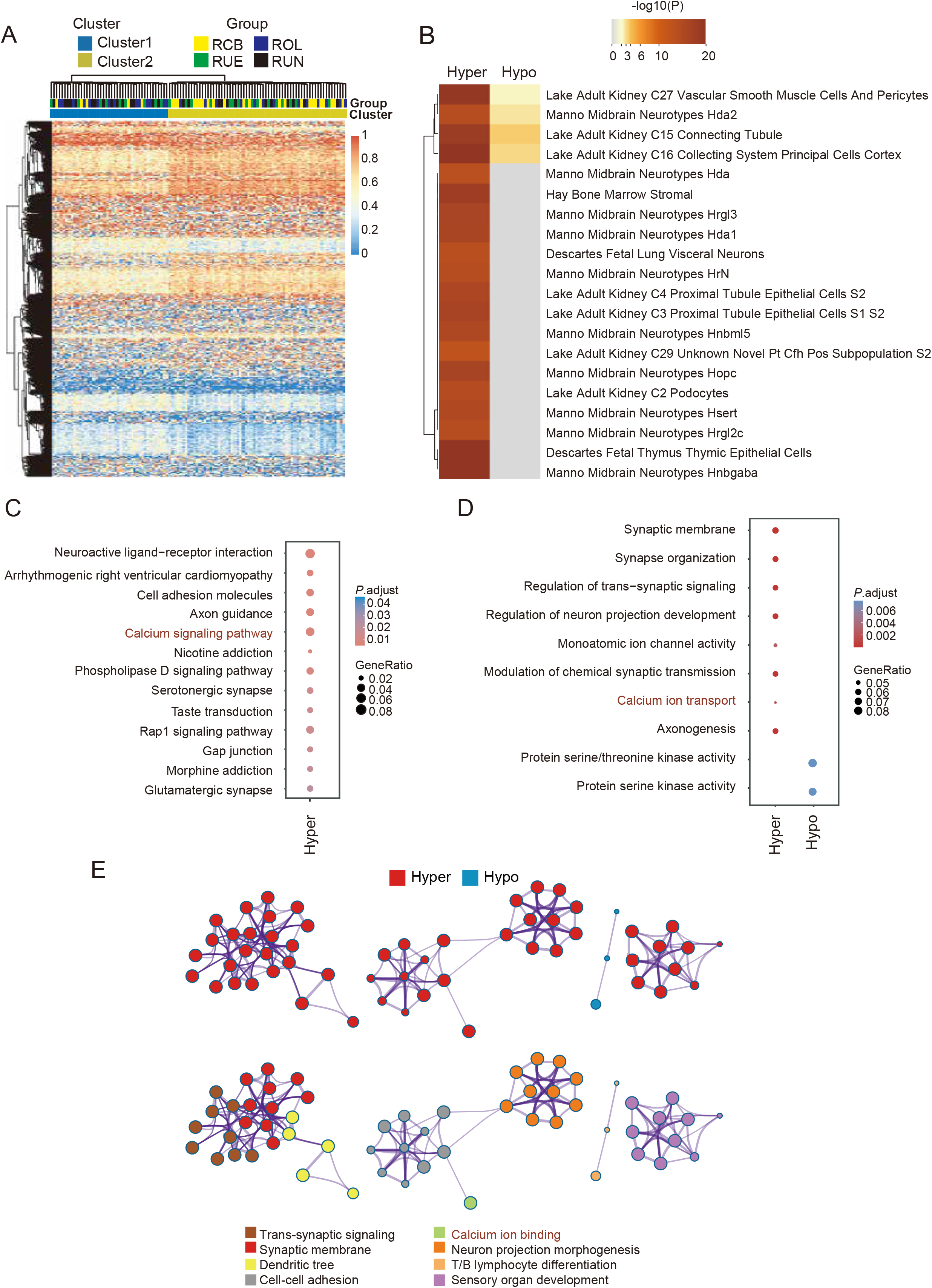
Annotation of DMSs between Cluster1 and Cluster2. (A) Heatmap plot showing the methylation value of the top 2000 DMSs between two clusters, with blue to red indicating increasing methylation levels (from low to high). Each column represents a sample, and each row represents a DMS. (B) Cell types enriched with differentially hyper and hypo DMS-annotated genes. (C) KEGG functional enrichment results, where only hyper DMS-annotated genes are enriched in pathways. (D) GO enrichment analysis of DMS-annotated genes. (E) The PPI network analysis of genes associated with differentially methylated sites derived by unsupervised clustering. DMSs, differentially methylated sites; KEGG, Kyoto encyclopedia of genes and genomes; GO, gene ontology.

Furthermore, Kyoto encyclopedia of genes and genomes (KEGG) and gene ontology (GO) enrichment analyses showed that the hypermethylated DMS-associated genes were both significantly enriched in the calcium-related biological processes and pathways, including calcium signaling pathway, calcium ion transport and calcium ion binding (**Fig. 3C–E**). Specifically, dysregulation of intracellular calcium homeostasis in macrophages can activate the *NLRP3* inflammasome, a key component of gout-related inflammation[26]. Moreover, calcium overload has been reported to impair insulin signaling in peripheral tissues, exacerbating insulin resistance, which in turn elevates renal *URAT1* expression and inhibits uric acid excretion [27]. KEGG and GO enrichment analyses additionally identified significant enrichment of cell adhesion molecules and the Rap1 signaling pathway, which have likewise been reported in differential DNA methylation analyses between gout patients and healthy individuals[20]. Notably, Rap1 has been shown to interact with calcium signaling in regulating platelet activation and neuronal plasticity[28], which is consistent with our observed enrichment of neuronal cell types (**Fig. 3B**) and the GO enrichment analysis (**Fig. 3D**). The PPI network of calcium ion transport and synaptic-related pathways further support the interplay between calcium signaling, synaptic plasticity, and neuronal viability (**Fig. 3E**). Furthermore, DNA methylation-based deconvolution revealed divergent immune microenvironments: Cluster 2 exhibited a significantly higher proportion of neutrophils, while Cluster 1 was enriched in memory B cells (**Fig. S3B)**. This observation is consistent with the enrichment of DMS-associated genes in T/B lymphocyte differentiation pathways, suggesting potential differences in immune microenvironment, inflammatory response, and disease progression between the two molecular subtypes.

Finally, among the KEGG pathways enriched by DMS-associated genes, the calcium signaling pathway contained significantly more overlapping genes between DMS-associated genes shared with GWAS-reported serum urate–related genes compared than random expectation (hypergeometric test, *P* ≤ 0.05, Supplementary Tables 6–8), i.e. *ERBB4*, *FGF5*, and *PDE1A*. The same analysis performed on GO enrichment identified pathways related to monoatomic ion channel complex and positive regulation of monoatomic ion transport, with major overlapping genes including *AKAP6*, *CLIC5*, *DPP6*, *KCNJ2*, *RELN*, and *STC1* (**Fig. S3C; Supplementary Table 6-8**). The concurrent enrichment of metal ion transport pathways influenced by DNA methylation is consistent with previous studies reporting that gout susceptible loci are enriched in vascular transport, transport across the blood-brain barrier[29]. Moreover, regulatory pathways involved in transmembrane ion transport have been shown to be closely related to serum urate concentrations and the pathogenesis of gout[30]. Collectively, our results further demonstrate that methylation alterations in these genes are closely associated with gout subtypes and serum urate levels and neural functional pathways.

### Methylation-based model achieves optimal performance in classifying gout patients with high and low FE_UA_

Given that the two patient clusters identified through unsupervised clustering were significantly associated with FE_UA_, suggesting the potential of methylation features as a surrogate for FE_UA_ in clinical applications, we further selected FE_UA_ as the classification label. Patients were stratified into high (FE_UA_ ≥ 5.5%) and low (FE_UA_ < 5.5%) FE_UA_ groups to construct classification models based on methylation profiles. To identify DNA methylation sites associated with FE_UA_-based classification, we first employed the least absolute shrinkage and selection operator (LASSO) regression method to select 18 methylation sites as key features at the minimum λ value (**Fig. S4A–B**). Based on these features, we built machine learning models, including LASSO, extreme gradient boosting (XGBoost), random forest (RF), and support vector machine (SVM) models to discriminate 108 gout patients. Additionally, to assess the robustness of feature selection, we constructed another LASSO classification model using the six methylation sites identified under the lambda.1se condition as a reference for comparison. The results showed that the XGBoost model achieved the best classification performance with an AUC of 0.961, significantly outperforming other models (**Fig. 4A**). The RF and SVM models also demonstrated strong classification capabilities, with AUC values of 0.922 and 0.934, respectively. We further applied the SHapley Additive exPlanations (SHAP) interpreter to rank feature importance in the XGBoost model to enhance interpretability, aiming to quantify the relative contribution of each methylation feature to FE_UA_ prediction and improve model interpretability (**Fig. S4C**). This feature importance ranking provided a critical basis for subsequent biological interpretation and functional exploration of the identified methylation loci. Notably, even when the number of features was reduced from 18 to the top 10 sites, the model retained high predictive accuracy with an AUC of 0.88 (**Fig. S4D**), indicating the stability and efficiency of the selected methylation markers. Besides, the 18 methylation sites were significantly correlated with multiple clinical characteristics (**Fig. 4B**), including serum creatinine, blood urea nitrogen, age, fasting glucose, UA, and FE_UA_, indicating the association between DNA methylation and phenotype.

**Figure 4.**
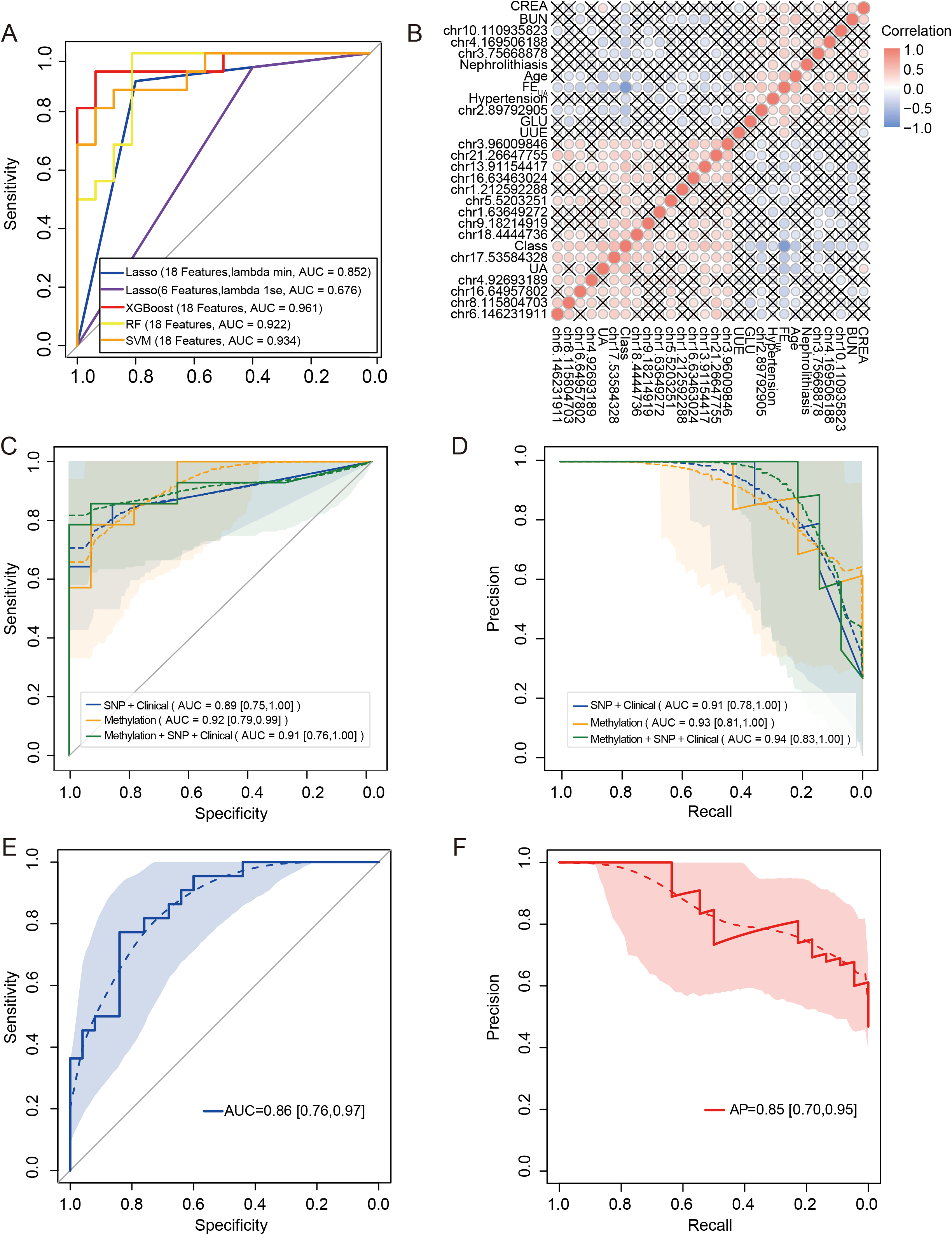
Classification model in classifying gout patients with high and low FE_UA_. (A) Comparative evaluation of ROC curves across four machine learning models developed using 18 methylation features, along with the LASSO classification model established using 6 methylation features selected under the LASSO regression lambda 1se condition, which served as a reference for comparison. (B) Association analysis between the 18 methylation features and clinical characteristics, with red indicating positive correlations, blue indicating negative correlations, and cross symbols denoting lack of statistical significance (*P* > 0.05). *P* values were calculated using the two sides test. ROC(C) curves and PR(D) curves of SGD classification models built on the same training set using SNP and clinical features, methylation features, and combined methylation, SNP, and clinical features, evaluated on the same test set. (E) ROC curve of the XGBoost-based classification model in the validation cohort. (F) AP curve of the XGBoost-based classification model in the validation cohort. LASSO, least absolute shrinkage and selection operator; XGBoost, extreme gradient boosting; RF, random forest; SVM, support vector machine; SGD, stochastic gradient descent. ROC, receiver operating characteristic; PR, precision-recall; AUC, area under the curve; AP, average precision; SNP, single-nucleotide polymorphism.

Furthermore, to enable a fair comparison of models constructed from different feature sets within the same test dataset, we rebuilt the previously reported model using six clinical variables and four single nucleotide polymorphism (SNP) genotypes based on an SGD algorithm[24]. The reproduced model achieved an AUC of 0.89 and an AP of 0.91, consistent with the reported results (**Fig. 4C–D**). In contrast, our model using 18 methylation features achieved an AUC of 0.92 and an AP of 0.93, while the integrated model combining methylation, SNP, and clinical features yielded an AUC of 0.91 and an AP of 0.94. Lastly, in an independent validation cohort of 100 individuals, the XGBoost model achieved an AUC of 0.86 and an AP of 0.85 (**Fig. 4E–F**), further confirming the predictive power and generalizability of methylation-based features. Collectively, these findings highlight the unique advantages and clinical potential of methylation signatures in predicting uric acid excretion capacity among gout patients.

### Potential clues of DNA methylation signature site-associated genes *NEK1* and *TRPS1* in the inflammatory process

We annotated the genomic positions and functional features of 18 methylation sites (**Supplementary Table 9).** Among them, 3 sites are in the promoter region, 11 in introns, and 4 in distal intergenic regions. Furthermore, epigenomic data annotation revealed that *NEK1*-associated methylation site chr4:169,506,188 is located within 1 kb downstream of the *NEK1* transcript (ENST00000638824.1) TSS and overlapped with a region of high H3K4me3 accessibility (**Fig. S5A**), suggesting its potential regulatory function. Motif analysis using the JASPAR database identified 11 potential transcription factor binding motifs with relative scores above 0.8 (**Supplementary Table 10**), with the top four corresponding to *ETS1*, *NFATC3*, *SPI1*, and *SPIB* (**Fig. S5B**). These results suggest that hypermethylation at chr4:169,506,188 in patients with high FE_UA_ may impede transcription factor binding or alter alternative splicing of *NEK1*, thereby reducing its expression.

To explore the expression of signature genes, we retrieved three publicly available single-cell RNA sequencing (scRNA-seq) datasets from peripheral blood mononuclear cells with available uric acid level. After quality control, cell-type annotation, and stratification based on uric acid levels (**Fig. S5C**), a total of 270, 041 cells were clustered to B cells, CD4 T cells, CD8 T cells, monocyte-macrophages (Mono-Macro), neutrophils, and natural killer (NK) cells (**Fig. 5A**), with classical marker genes shown in **Fig. S5D**. Expression profiling of genes annotated to the 18 methylation sites revealed that *NEK1* (chr4:169,506,188) and *TRPS1* (chr8:115,804,703) displayed high expression across multiple immune cell types and had opposite methylation trends between the FE_UA_-High and FE_UA_-Low subgroups (**Fig. 5B-E and Fig. S5E-F**). *NEK1* is predominantly expressed in T, B and NK cells (**Fig. 5D**), suggesting influenced lymphocyte processes via DNA damage response, cell cycle, and apoptosis [31], with its expression correlates with immune cell markers [32]. Additionally, *NEK1* variants linked to cystic kidney disease and nephrolithiasis [33] and its overexpression in mice (*mNek1*) suppresses ciliogenesis and affects renal epithelial cell function[34], suggesting its role in renal function. In contrast, *TRPS1* showed higher expression in macrophages, monocytes, and neutrophils (**Fig. 5E**), suggesting its potential involvement in regulating myeloid cell inflammatory phenotypes. The significant expression differences of *NEK1* and *TRPS1* across UA-stratified patient groups (**Fig. 5B**, **Fig. 5C**, and **Fig. S5F**) further supporting their association with uric acid excretion phenotypes and immune-inflammatory processes.

**Figure 5.**
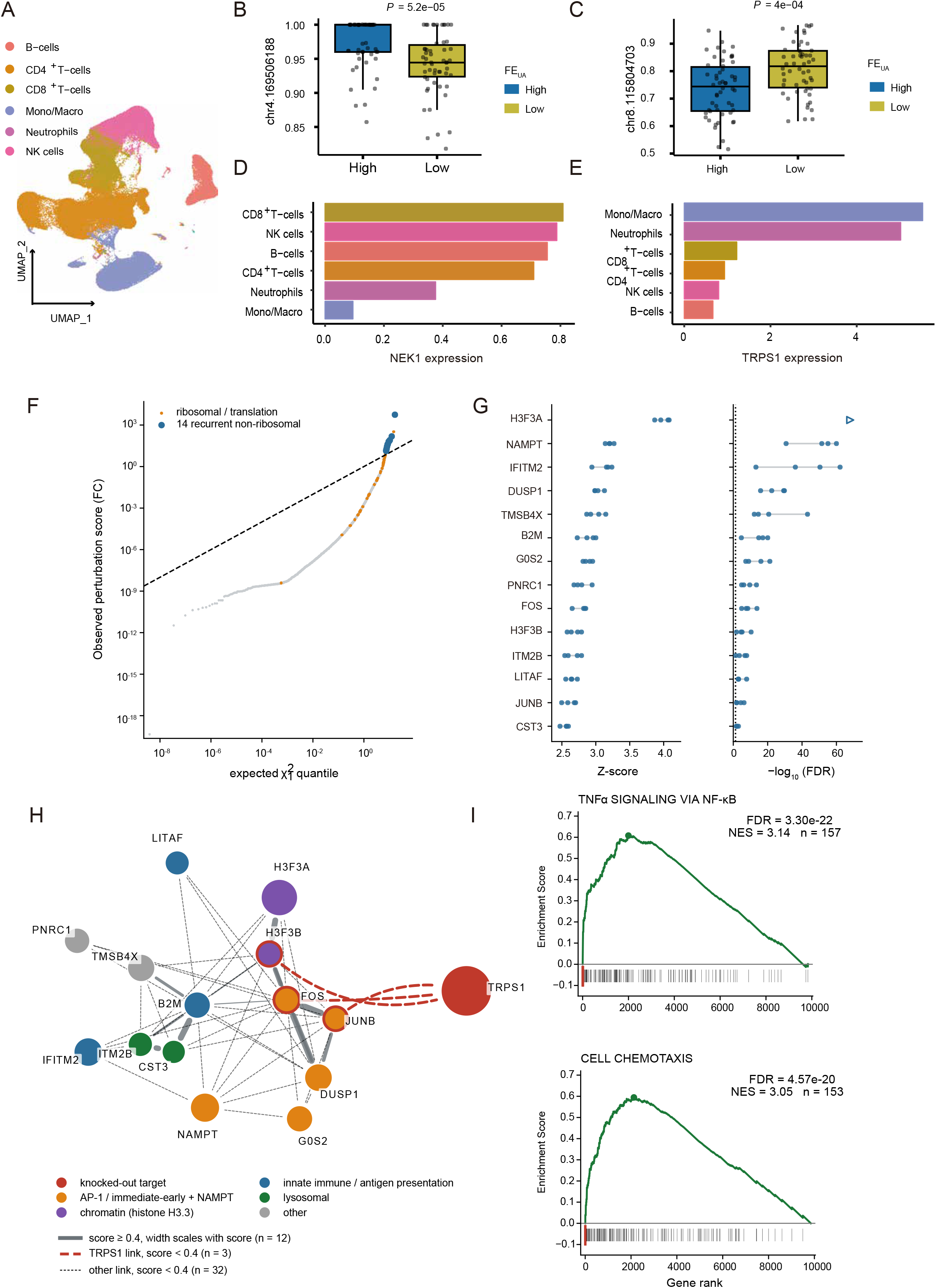
Methylation-associated candidate genes in gout single-cell transcriptomes and in silico knockout of *TRPS1*. (A) UMAP dimensionality reduction to visualize the annotated cell populations. Three scRNA-seq datasets (GSE211783, GSE217561, PRJNA861849) derived from the peripheral blood of 19 gout patients. Differential DNA methylation analysis of the locus chr4:169,506,188 (B) and chr8:115,804,703 (C) between FE_UA_-High and FE_UA_-Low subgroups. Average expression of the associated gene *NEK1* (D) and *TRPS1* (E) across different cell types. (F) Q–Q plot of the scTenifoldKnk perturbation score (FC) against the χ²□ null for the 10,082 tested genes, excluding *TRPS1* (seed 1); dashed line, y = x; both axes log. Orange, ribosomal/translation genes; blue, the 14 non-ribosomal genes significant in all four seeds; grey, remainder. (G) Z score (left) and −log□□ (FDR) (right) for those 14 genes, ordered by mean rank. Four points per gene, one per seed (1, 2, 3, 42); grey bar, range. Dotted line, FDR = 0.05. Open triangle, *H3F3A* (FDR underflowed to 0 in all runs). (H) STRING association network of the 14 genes and *TRPS1*. Node area scales with mean knockout Z score; colour, functional module. Grey solid edges, high-to-medium confidence, width scaling with score; red dashed, the sub-threshold associations involving *TRPS1* (partner and score labelled); black dashed, the remaining sub-threshold associations. Red rings, *TRPS1* partners. (I) GSEA running enrichment score for the two top-ranked gene sets, on the ranking from which ribosomal/translation genes were removed. Black ticks, set members; red ticks, the 14 recurrent genes. UMAP, uniform manifold approximation and projection; scRNA-seq, single-cell RNA sequencing; FE_UA_, fractional excretion of uric acid; FC, perturbation score; FDR, false discovery rate; NES, normalized enrichment score.

Given the myeloid-restricted expression of *TRPS1*, we performed an in silico knockout of T*RPS1* in monocyte-macrophages to identify its downstream programme. 14 significantly differentially expressed genes were recovered reproducibly across random seeds **(Fig. 5F, G**): *H3F3A*, *NAMPT*, *IFITM2*, *DUSP1*, *TMSB4X*, *B2M*, *G0S2*, *PNRC1*, *FOS*, *H3F3B*, *ITM2B*, *LITAF*, *JUNB* and *CST3*. *TRPS1* had no high-confidence STRING association with any of them, as expected for a DNA-binding repressor acting without a physical complex, whereas the 14 genes themselves formed a connected network built around an AP-1 core (FOS–JUNB–DUSP1) with B2M as its hub (**Fig. 5H**). Gene set enrichment analysis placed TNFα signaling via NF-κB first (NES = 3.14, FDR = 3.3 × 10□ ²²; **Fig. 5I**), with 7 of the 14 genes in its leading edge. The parallel knockout of *NEK1* in lymphoid populations recovered no comparable module. These results suggest that *TRPS1*, itself under differential methylation control between FE_UA_ subgroups, sustains an AP-1/NF-κB inflammatory programme in monocyte-macrophages.

Collectively, the distinct expression patterns of *NEK1* and *TRPS1* across immune cell types and UA subgroups, together with evidence of AP-1/NF-κB inflammatory programme in silico, providing an clue of potential regulatory associations between DNA methylation and gout pathogenesis.

## Discussion

In this study, we comprehensively analyzed genome-wide DNA methylation profiles from 208 gout patients across four clinical subtypes. By exploring methylation heterogeneity via clinically defined subgroups and unsupervised clustering, we characterized differential methylation patterns, functional pathways, cell type enrichments, and PPI networks. We found gout methylation heterogeneity was primarily associated with FE_UA_, highlighting a strong link between epigenetic signatures and urate excretion capacity. Based on these features, we constructed classification models tailored to various diagnostic scenarios. The FE_UA_ model achieved optimal performance, demonstrating the potential of methylation-based gout stratification, and providing new insights into the biological basis of gout heterogeneity.

Currently, studies on gout classification are mainly based on uric acid excretion indices (e.g., FE_UA_, UUE) and other comorbidities. In our cohort, the RCB subtype was associated with a younger age at disease onset, higher SU levels, and higher eGFR, whereas RUN associated with older age at onset, lower SU levels, and reduced eGFR (Table S1 and S2). Although similar non-significant trends occurred in RUE and ROL subtypes, likely due to sample size constraints. Beyond clinically defined classifications, previous studies have applied clustering approaches using heterogeneous sources of information, including comorbidity profiles, clinical characteristics, and biological markers, identifying three to five gout subtypes across populations[35–37]. However, these depend highly on the specific data modalities used. In contrast, our DNA methylation-based framework introduces a stable, complementary molecular omics dimension rarely explored in gout research. By capturing epigenetic alterations that integrate genetic susceptibility and environmental influences, this approach provides a complementary and potentially more stable stratification beyond conventional clinical phenotypes. Although detailed comorbidity data were unavailable, associations between methylation-defined subtypes, comorbidities, and clinical outcomes (e.g., drug response, flare frequency) warrant future investigation. Notably, this molecular classification may have implications for drug response, flare frequency, and gout-related complications.

Biologically, membrane potential–related pathways were significantly enriched in the RUN subtype. RUN patients typically exhibit preserved renal urate excretion, suggesting their hyperuricemia stems from impaired intestinal urate metabolism[8]. Several RUN-associated methylation sites were annotated to solute carrier family genes (Fig 1E), which play key roles in regulating urate transport and organic anion homeostasis[38, 39]. These findings highlight potential therapeutic targets for the RUN subtype, for which specific treatments are currently lacking. Additionally, intersection GWAS signals with enriched pathways identified shared genes, including *ERBB4*, *FGF5*, *PDE1A*, *AKAP6*, *CLIC5*, *DPP6*, *KCNJ2*, *RELN*, and *STC1*. Among these genes, *STC1* has been repeatedly implicated in serum urate regulation in trans-ethnic GWAS meta-analyses[40]. Notably, SNPs in CLIC5 and KCNJ2 are linked to RUN and ROL subtypes, respectively[41]. Consistently, both *CLIC5* and *KCNJ2* were significantly associated with Cluster1 in our methylation-based classification, which overlaps with RUN and ROL phenotypes. Together, these results demonstrate concordance between genetic associations and epigenetic subtyping, supporting a functional role for these genes in urate homeostasis and gout pathophysiology, even for loci not reaching genome-wide significance[42, 43].

Translationally, our 18-CpG methylation-based FE_UA_ classification model outperformed previous clinical and SNP-based models[24]. Unlike SNP models focusing mainly on urate variants, our model incorporates CpG sites linked to urate metabolism and immune-inflammatory pathways, offering comprehensive stratification and target discovery. Furthermore, increasingly accessible PCR-based assays offer a noninvasive method for gout stratification, potentially enhancing personalized care[44]. Nevertheless, the observed decline in AUC and average precision in an independent validation cohort indicates that the generalizability of the model requires further optimization and validation in larger and more heterogeneous populations. Integrating scRNA-seq data, we identified *NEK1* and *TRPS1* as potential genes under epigenetic regulation associated with gout subtypes, uric acid levels and immune-inflammatory processes. *NEK1*, a multifunctional serine/threonine kinase, has been shown to regulate glucose metabolism by modulating the membrane localization of glucose transporter 1 (*GLUT1*), as evidenced by reduced *GLUT1* membrane expression following *NEK1* depletion in mouse embryonic fibroblasts[45, 46]. Notably*, GLUT1* is upregulated and enriched on the macrophage membrane upon MSU crystal stimulation, enhancing glycolytic activity to support the assembly and activation of the *NLRP3* inflammasome and subsequent interleukin-1β (IL-1β) maturation and secretion. Inhibition of *GLUT1*, either by glucose deprivation or pharmacological blockade (e.g., STF-31), markedly attenuates MSU-induced *NLRP3* activation, IL-1β release, and inflammatory responses[47]. Therefore, that the methylation of *NEK1* may play a regulatory role in gout-related inflammation by modulating *NLRP3* inflammasome activation and cytokine maturation. Similarly, *TRPS1* encodes a zinc-finger transcription factor that may modulate inflammatory responses through inhibition of STAT signaling or modulation of NF-κB activity in myeloid cells. Experimental evidence indicates that *TRPS1* can directly repress *STAT3* expression[48], and given the close cooperation between *STAT3* and NF-κB in inflammatory signaling networks[49], this provides a plausible mechanistic basis for *TRPS1*-mediated regulation of inflammation. This is supported by in silico knockout of *TRPS1*.

Despite these promising findings, several limitations remain. The current methylation dataset of 208 gout samples, although representative, remains relatively small and may not fully capture the diversity across subtypes. Additionally, models were developed in a Chinese male cohort require validation in larger, multi-ethnic populations is required to ensure generalizability. Future studies should establish multi-center cohorts and develop cost-effective targeted methylation panels to improve clinical applicability. Moreover, integrative multi-omics analyses, including single-cell methylome and transcriptome profiling, are needed to better elucidate the molecular mechanisms and cellular origins underlying gout heterogeneity. Finally, experimental validation of *NEK1* and *TRPS1* methylation levels across diverse biological conditions is warranted to confirm their functional contributions to gout subtype differentiation and disease progression.

## Conclusions

In conclusion, this study provides the first comprehensive methylome-based molecular subtyping of gout, revealing distinct epigenetic signatures associated with urate excretion phenotypes and pathways. By linking molecular epigenetic profiles to clinical phenotypes of urate metabolism, we provide new insights into the biological basis of gout heterogeneity. Our results highlight the potential of DNA methylation as both a biomarker and a mechanistic link bridging genetic susceptibility, metabolic dysregulation, and clinical heterogeneity in gout. Moving forward, integrating multi-omics data and expanding validation across diverse populations will further refine these molecular subtypes and accelerate the translation of epigenetic biomarkers into precision diagnosis and targeted therapy for gout.

## List of abbreviations

ACR/EULAR: American College of Rheumatology/European League Against Rheumatism;
AP: average precision
AP-1: activator protein 1;
AUC: area under the curve;
CD: cluster of differentiation;
CpG: cytosine-phosphate-guanine dinucleotide
DMS: differentially methylated site;
DNA: deoxyribonucleic acid;
eGFR: estimated glomerular filtration rate;
FC: fold change relative to background;
FDR: false discovery rate;
FE_UA_: fractional excretion of uric acid
gDNA: genomic DNA;
GLUT1: glucose transporter 1
GO: Gene Ontology;
GSA: Genome Sequence Archive
GSEA: gene set enrichment analysis
GWAS: genome-wide association study
IL-1β: interleukin-1 beta;
KEGG: Kyoto Encyclopedia of Genes and Genomes
LASSO: least absolute shrinkage and selection operator
MSigDB: Molecular Signatures Database;
MSU: monosodium urate
NEK1: NIMA-related kinase 1;
NES: normalized enrichment score
NF-κB: nuclear factor kappa B
NK: natural killer;
NLRP3: NLR family pyrin domain containing 3
PBMC: peripheral blood mononuclear cell;
PCA: principal component analysis
PCR: polymerase chain reaction
PPI: protein-protein interaction;
PR: precision-recall
Q-Q: quantile-quantile
RCB: renal urate combined;
RF: random forest;
ROC: receiver operating characteristic
ROL: renal urate overload;
RUE: renal urate underexcretion;
RUN: renal urate normal
scRNA-seq: single-cell RNA sequencing
sCr: serum creatinine;
SGD: stochastic gradient descent
SHAP: SHapley Additive exPlanations
SNP: single-nucleotide polymorphism;
STAT3: signal transducer and activator of transcription 3;
STRING: Search Tool for the Retrieval of Interacting Genes/Proteins
SU: serum urate;
sUA: serum uric acid;
SVM: support vector machine
TNF-α: tumor necrosis factor alpha;
TRPS1: transcriptional repressor GATA binding 1
TSS: transcription start site;
UA: uric acid;
uCr: urine creatinine;
ULT: urate-lowering therapy;
UMAP: uniform manifold approximation and projection
URAT1: urate transporter 1;
uUA: urinary uric acid;
UUE: 24-hour urinary uric acid excretion
vKO: virtual knockout;
WGBS: whole-genome bisulfite sequencing
XGBoost: extreme gradient boosting.

## Declarations

### Ethics approval and consent to participate

This study was approved by the Ethics Committee of the Affiliated Hospital of Qingdao University (approval number: QYFYKYLL923011921). All participants gave their written informed consents.

### Consent for publication

All authors had full access to the data, approved the final manuscript, and accept responsibility for the decision to submit for publication.

### Availability of data and materials

The raw whole-genome bisulfite sequencing (WGBS) data have been deposited in the Genome Sequence Archive (GSA) under the National Genomics Data Center and the National Center for Bioinformation / Beijing Institute of Genomics, Chinese Academy of Sciences (GSA-Human: HRA010464 and HRA013297; https://ngdc.cncb.ac.cn/gsa-human) [50]. Other data supporting the findings of this study are available from the corresponding author upon reasonable request.

### Competing interests

The authors declare no competing interests.

### Fundings

Strategic Priority Research Program of the Chinese Academy of Sciences (XDA0460403), the National Natural Science Foundation of China (82220108015), and the National Key Research and Development Program of China (2022YFC2503304).

### Author contributions

Guangmin Zheng: Methodology, Software, Data curation, Formal analysis, Visualization, Conceptualization, Writing - original draft, Writing - review & editing. Aichang Ji: Data curation, Investigation, Validation, Writing - review & editing.

Wenrui Shi: Investigation, Validation, Writing - review & editing.

Xinde Li: Investigation, Validation, Writing - review & editing.

Xiuyan Ruan: Investigation, Validation, Writing - review & editing.

Xiaopeng Ji: Investigation, Validation, Writing - review & editing.

Xiangdong Fang: Supervision, Funding acquisition, Project administration, Resources, Writing - review & editing

Changgui Li: Supervision, Funding acquisition, Resources, Writing - review & editing Hongzhu Qu: Supervision, Conceptualization, Funding acquisition, Visualization, Project administration, Resources, Writing - original draft, Writing - review & editing

## Acknowledgements

We would like to thank the hospital staff for their help with cohort recruitment and sample collection. And we are also grateful to the patients and volunteers who provided the samples for our study.

During the preparation of this manuscript, the authors used ChatGPT / Gemini to check spelling and grammar and assist with literature search strategies. After using this tool, the authors reviewed and edited the content as needed and take full responsibility for the content of the publication.

## Notes

https://ngdc.cncb.ac.cn/search/specific?db=hra&q=HRA010464

https://ngdc.cncb.ac.cn/search/specific?db=hra&q=HRA013297

### Competing Interest Statement

The authors have declared no competing interest.

