## SupplementaryMaterial for "DNA methylation reveals molecular heterogeneity and urate excretion-associated subtypes of gout"

**This PDF file includes:**

Supplementary methods

References

Supplementary figure legends

### **Supplementary methods**

#### **WGBS data preprocessing**

The preprocessing of WGBS data primarily included raw data quality control, alignment, and evaluation of capture efficiency and sequencing depth. Raw reads generated by the sequencer contained adapters and low-quality bases, which could affect downstream assembly and analysis. To obtain high-quality clean reads, fastp v0.23.4[1] was used for filtering. Briefly, reads containing adapters, reads with more than 10% unknown nucleotides, and reads in which more than 50% of bases had low quality scores (Q ≤ 30) were removed.

Cleaned FASTQ data were aligned to the hg38 reference genome. For each sample, sequencing data were processed using the BatMeth2 pipeline[2] in an automated workflow. Samtools v1.19.2[3] was employed for downstream processing. The pipeline generated tab-delimited output files containing information such as CpG positions, methylated cytosine read counts at each position, total CpG coverage, and methylation distributions in CH, CHG, and CHH contexts. Genome-wide methylation levels were calculated as the percentage of methylated cytosines for each sequence context (CG, CHG, CHH) across the whole genome, individual chromosomes, and specific genomic regions. To evaluate methylation patterns in different genomic regions, methylation profiles were plotted for 2-kb flanking regions and gene bodies (or transposable elements) based on the average methylation level within each window. A custom bash script was used to merge approximately 400,000 methylated cytosine sites per individual into a combined methylation matrix across all samples, formatted for compatibility with the methylKit v1.28.0 R package [4] for subsequent analyses.

#### **WGBS methylation analysis**

Unsupervised clustering analysis was performed using methylKit (v1.28.0) [4] in the R environment (v4.3.1). Methylation data were first imported with the methRead function, normalized using normalizeCoverage function, and integrated with the unite function. Sample clustering was then conducted with the clusterSamples function, specifying dis = "correlation", method = "ward", and plot = TRUE. The resulting hierarchical clustering dendrogram was used to evaluate the global methylation similarity among samples. To determine the optimal number of clusters, the cascadeKM function from the vegan R package was applied [5]. Differential methylation analysis was carried out using the calculateDiffMeth function in methylKit, with family history, age, and other relevant covariates included as adjustment factors. Differentially methylated positions (DMPs) were filtered using the getMethylDiff function, applying a cutoff of methylation difference ≥10% and q-value ≤0.05. Sites were classified as hyper- or hypomethylated based on the meth.diff column in the result table.

Genomic annotation of DMPs was performed using the genomation (v1.34.0) package [6].Gene-level annotation was conducted with the annotatePeak function in ChIPseeker (v1.38.0) [7], with the tssRegion parameter set to ±3,000 bp. Genes annotated to DMPs were subjected to functional enrichment analysis using the enrichKEGG and enrichGO functions in clusterProfiler v4.10.1 [8]. Enrichment results were visualized with ggplot2 (v3.5.1) [9] and enrichplot (v1.22.0). Broader enrichment and network-based analyses, including cell type enrichment and protein–protein interaction (PPI) network analysis, were performed on the Metascape platform (v3.5.20240901) [10]. The resulting networks were rendered and visualized in Cytoscape (v3.8.2) [11].

#### **Estimation of cell type composition**

Considering that cell type composition is an important confounding factor in DNA methylation analysis of blood samples, we estimated the proportions of various leukocyte subtypes using the EpiDISH package (v2.18.0) [12]. Reference CpG probes associated with specific cell types, as summarized in previous studies, and implemented in EpiDISH, were utilized. These reference data were derived from Salas et al. and included 12 blood cell subtypes: CD8 naive T cells, CD8 memory T cells, CD4 naive T cells, CD4 memory T cells, regulatory T cells (Tregs), natural killer (NK) cells, naive B cells, memory B cells, monocytes, neutrophils, basophils, and eosinophils [13]. The epidish function in EpiDISH was employed to perform deconvolution-based estimation of cell type proportions.

#### **Feature selection and classification model construction**

For the clinical four-class classification model, methylation features from the discovery cohort were used as input to construct models. The dataset was randomly divided into training and testing sets at a 7:3 ratio. During model training, hyperparameter tuning was performed using grid search combined with five-fold cross-validation based on random forest (RF), extreme gradient boosting (XGBoost), and stochastic gradient descent (SGD), and models were evaluated on the testing set.

For the binary classification model, methylation features from the discovery cohort were used as input. The dataset was similarly split into training and testing sets in a 7:3 ratio. In the training set, feature selection and internal validation were performed using least absolute shrinkage and selection operator (LASSO) regression combined with 10-fold cross-validation[14] . Predictive covariates were selected based on the minimum λ criterion (lambda.min) using the R package glmnet (v4.1.8). The variables identified by LASSO were incorporated into machine learning models to construct classifiers for the target classification. Multiple machine learning algorithms were compared, including LASSO, support vector machine (SVM), RF, SGD, decision tree, and XGBoost. To validate classifier performance, the models were evaluated on the testing set and compared with previously established models, and the best-performing models were evaluated on the validation cohort. Model performance was assessed using receiver operating characteristic (ROC) curves and precision–recall curves (PRC). All modeling and analysis were conducted in R (v4.3.1) and Python (v3.8.3).

#### **scRNA-seq data acquisition, processing, and cell type annotation**

To investigate differences in RNA expression levels of genes corresponding to key methylation sites identified in the models across gout patients with varying serum urate levels, three publicly available gout-related scRNA-seq datasets, GSE211783[15], GSE217561[16], and PRJNA861849[17], were retrieved, comprising a total of 19 gout patient samples. The raw sequencing data from three datasets were downloaded and processed using a standardized pipeline. Sequencing reads from the scRNA-seq libraries were aligned to the human reference genome (GRCh38) using Cell Ranger v7.0.0 (10x Genomics). Data processing and downstream analysis were performed with the Seurat package v4.4.0 in R [18]. Cells with fewer than 200 detected genes or mitochondrial gene content greater than 20% were removed. Doublets were identified and removed using the DoubletFinder package v2.0.3. Batch effects among datasets were corrected using the CCA algorithm implemented in Seurat. Cell type annotation was conducted using the BlueprintEncode reference dataset with the SingleR package v2.0.0 [19]. Sample information and grouping details are provided in **Supplementary Table 3**. The number of cells obtained after unified quality control for each sample is summarized in **Supplementary Table 4**. Cluster-specific marker genes identified after annotation are listed in **Supplementary Table 5**.

### **Supplementary figure legends**

**Figure S1.** **Performance evaluation of DNA methylation–based models for four clinical gout subtypes.** (A) ROC and PR curves of the RF, XGBoost, and SGD models constructed for the clinical four-class classification task in the test dataset. (B) Venn diagram showing the overlap of DMSs among the four clinical gout subtypes. (C) Gene annotation and functional enrichment analysis of DMSs across four distinct gout subtypes. Blue lines indicate shared genes annotated from DMSs, and purple lines represent enriched common functional pathways. ROC, receiver operating characteristic; PR, precision-recall; AUC, area under the curve; AP, average precision; RF, random forest; XGBoost, extreme gradient boosting; SGD, stochastic gradient descent; DMSs, differentially methylated sites. RCB, renal urate combined; RUE, renal urate underexcretion; ROL, renal urate overload; RUN, renal urate normal.

**Figure S2.** **Unsupervised clustering analysis of gout patients in the validation cohort. (A) Correlation analysis of DNA methylation features of the discovery cohort.** (B) Determination of the optimal number of clusters. The red dot indicates the optimal cluster number. (C) Unsupervised clustering dendrogram in the validation cohort. (D) Correlation analysis between the unsupervised clustering results and clinical subtypes in the validation cohort, assessed by *Fisher’s* exact test. (E) Comparative analysis of FE_UA_ levels between the two clusters identified by unsupervised clustering in the validation cohort. *P* values were calculated using the *Wilcoxon* rank sum test. FE_UA_, the fractional excretion of uric acid.

**Figure S3.** **Key gene-pathway and enrichment network of DMS-annotated genes between Cluster1 and Cluster2.** (A) Chromosomal distribution of DMSs. (B) The composition of B memory cells (left) and neutrophils (right) showed significant differences between Cluster1 and Cluster2. *P* value was calculated with *Wilcoxon* test. **P* < 0.05. (C) Genes significantly regulating pathways related to positive regulation of monoatomic ion transport, regulation of metal ion transport, regulation of monoatomic ion transmembrane transport, and monoatomic ion channel complex in the GO pathway enrichment results, genes highlighted in red indicate those overlapping with GWAS-identified genes. FE_UA_, the fractional excretion of uric acid; DMSs, differentially methylated sites; KEGG, Kyoto encyclopedia of genes and genomes; PPI, protein-protein interaction. GO, gene ontology.

**Figure S4.** **Feature selection and feature importance ranking.** (A) LASSO regression for feature selection with determination of the optimal regularization parameter λmin via cross-validation. (B) The 18 methylation sites selected under λmin. (C) Feature importance ranking of the XGBoost-based classification model, interpreted using the SHAP explainer. (D) ROC curves of three classification models reconstructed using the top 10 SHAP-ranked features. XGBoost, extreme gradient boosting; SHAP, Shapley additive explanations; ROC, receiver operating characteristic; PR, precision-recall; AUC, area under the curve.

**Figure S5.** **Integration of single-cell RNA sequencing data and genomic context of gout-associated methylation sites.** (A) Genomic features within 500 bp upstream and downstream of the Chr4:169,506,188 were visualized in the UCSC Genome Browser. Synchronously displayed tracks include multiple transcript variants of NEK1, expression profiles, CpG island annotations, chromatin accessibility levels, and ChIP-seq binding signals for TFs (ETS1, NFATC3, RELB, and SPI1). (B) Potential binding motifs for the Chr4.169506188 methylation site, with the top 4 ranked by relative score. (C) UMAP visualization of integrated peripheral blood scRNA-seq data from gout patients after batch correction, showing well-aligned cellular distributions across datasets. (D) Marker gene expression patterns used for cell type annotation. (E) Differentially expressed genes associated with classification models methylation sites in scRNA-seq of gout patients across distinct cell types. (F) Differentially expressed genes associated with classification models methylation sites in scRNA-seq of gout patients grouped by UA, with non-gout individuals serving as the normal control group.UMAP, uniform manifold approximation and projection; scRNA-seq, single-cell RNA sequencing; UCSC, university of California, Santa Cruz; ChIP-seq, chromatin immunoprecipitation sequencing; TFs, transcription factor.
