## Supplementary figures and images for "DNA methylation reveals molecular heterogeneity and urate excretion-associated subtypes of gout"

### Figure S1

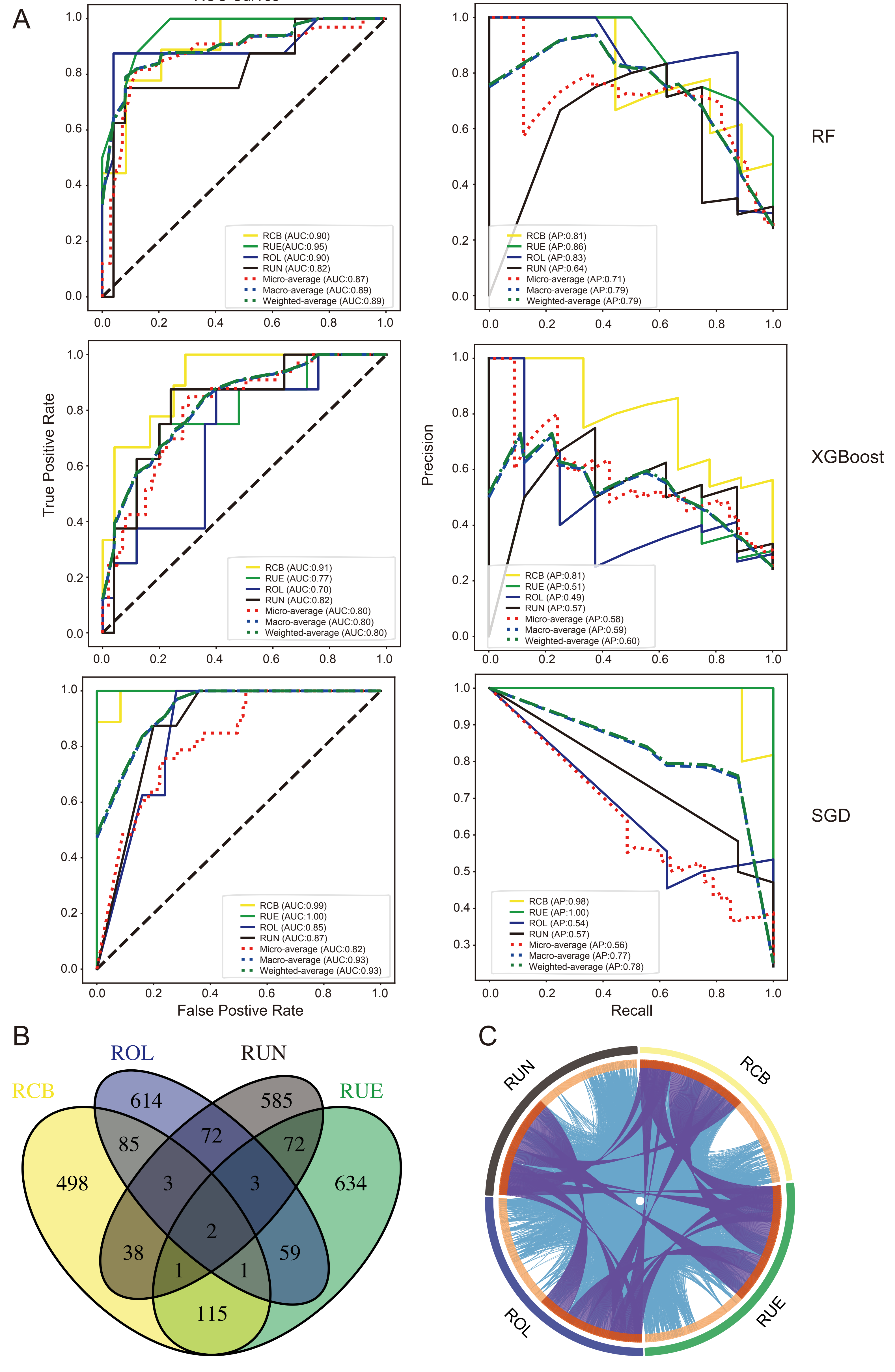

### Figure S2

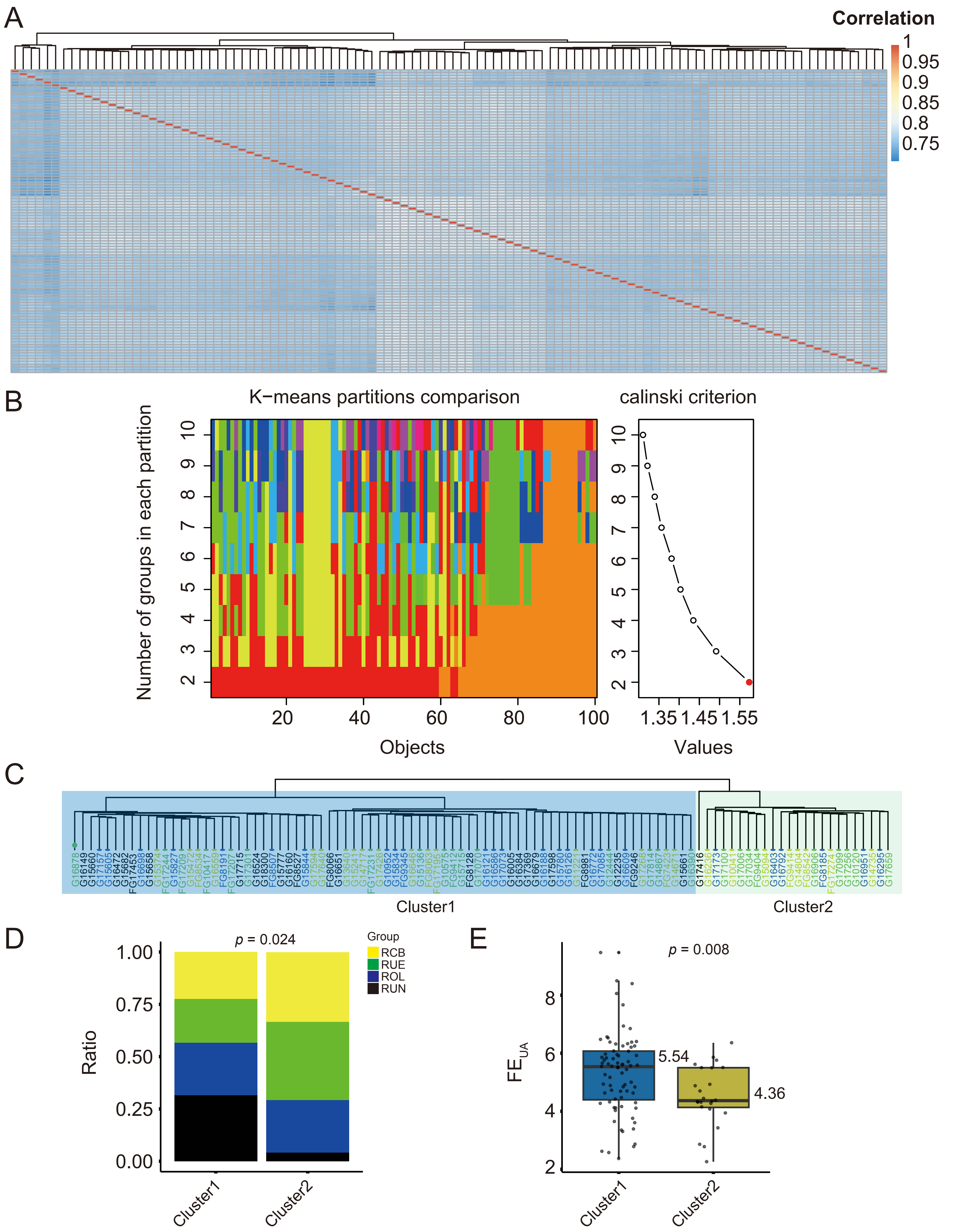

### Figure S3

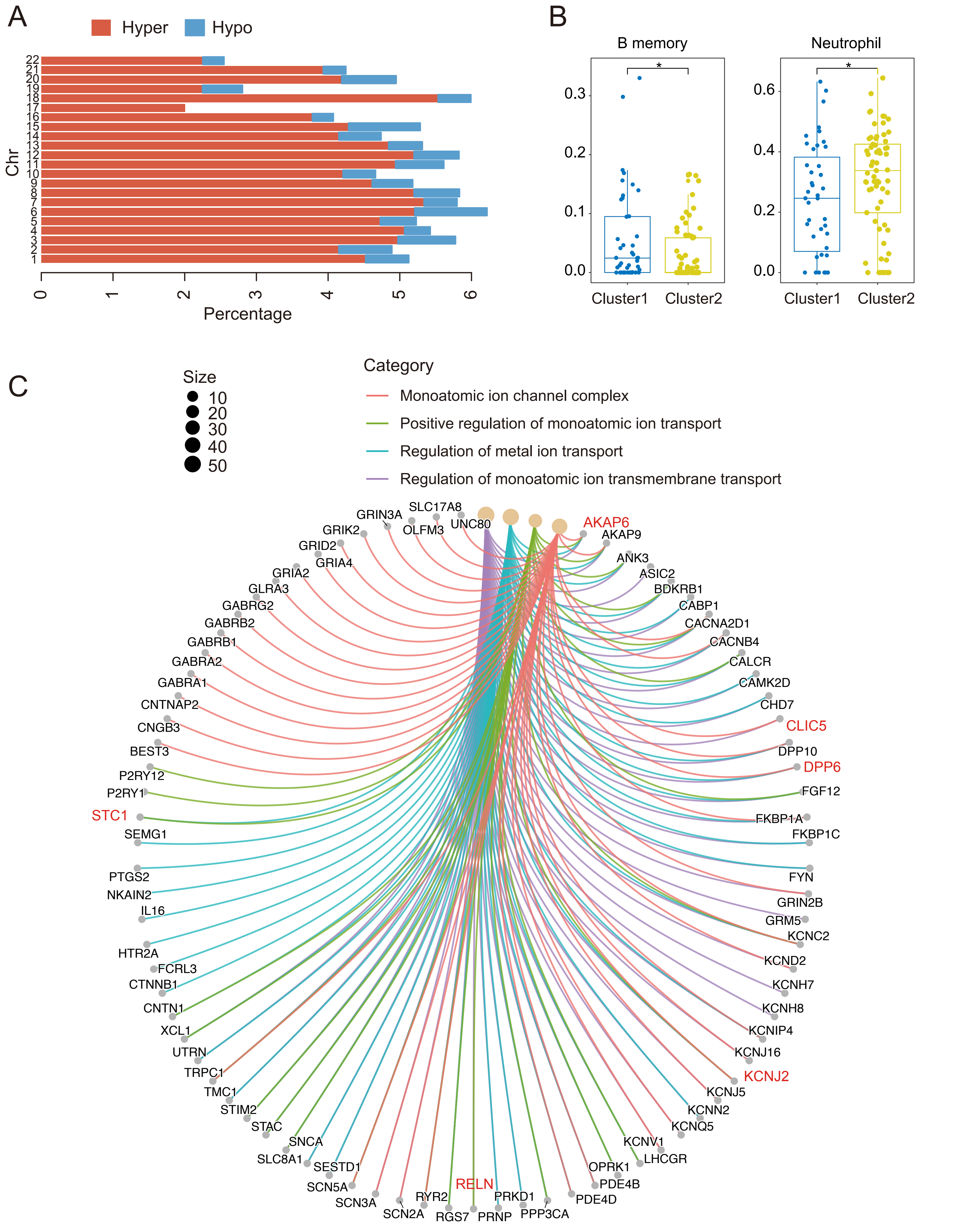

### Figure S4

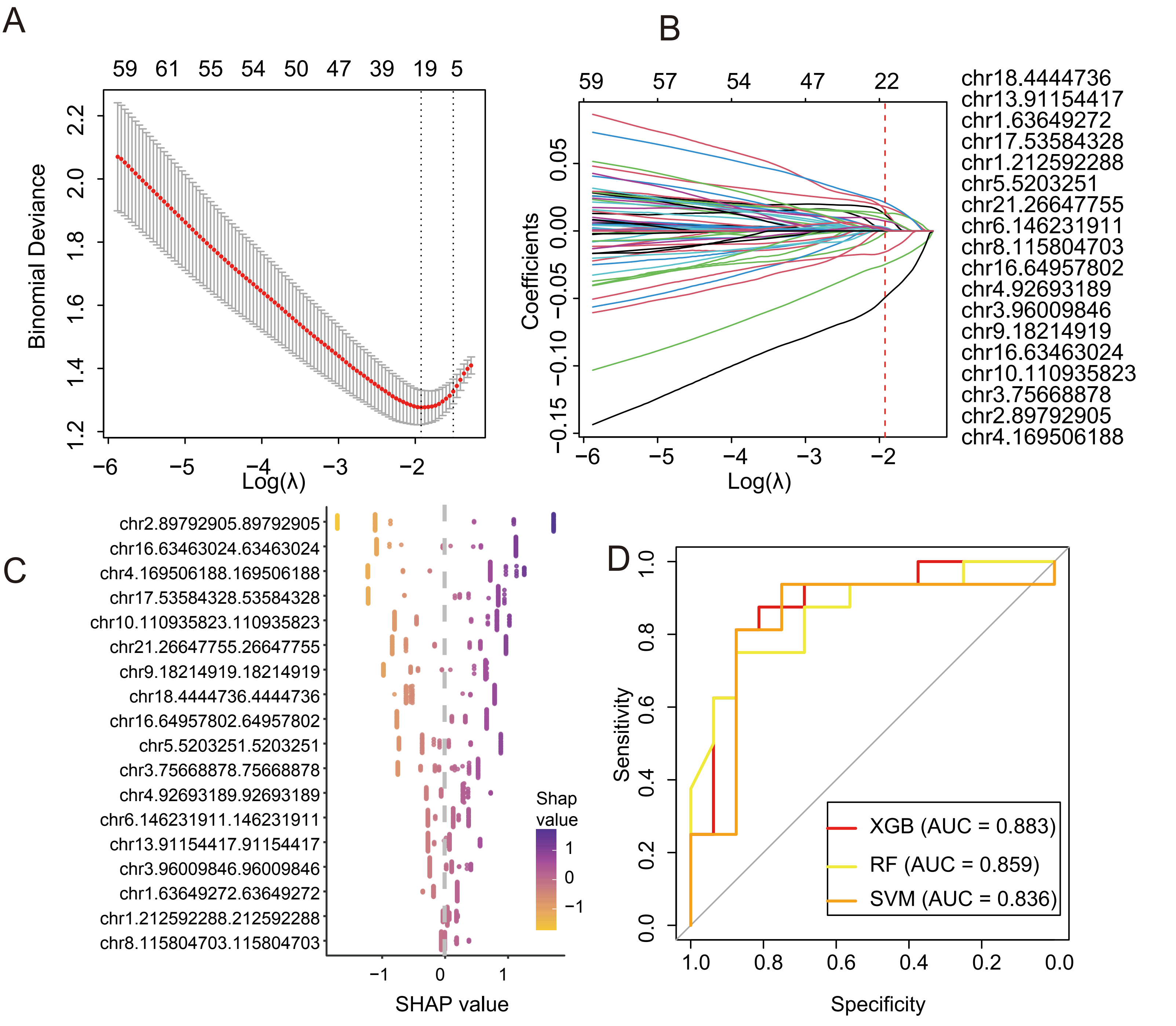

### Figure S5

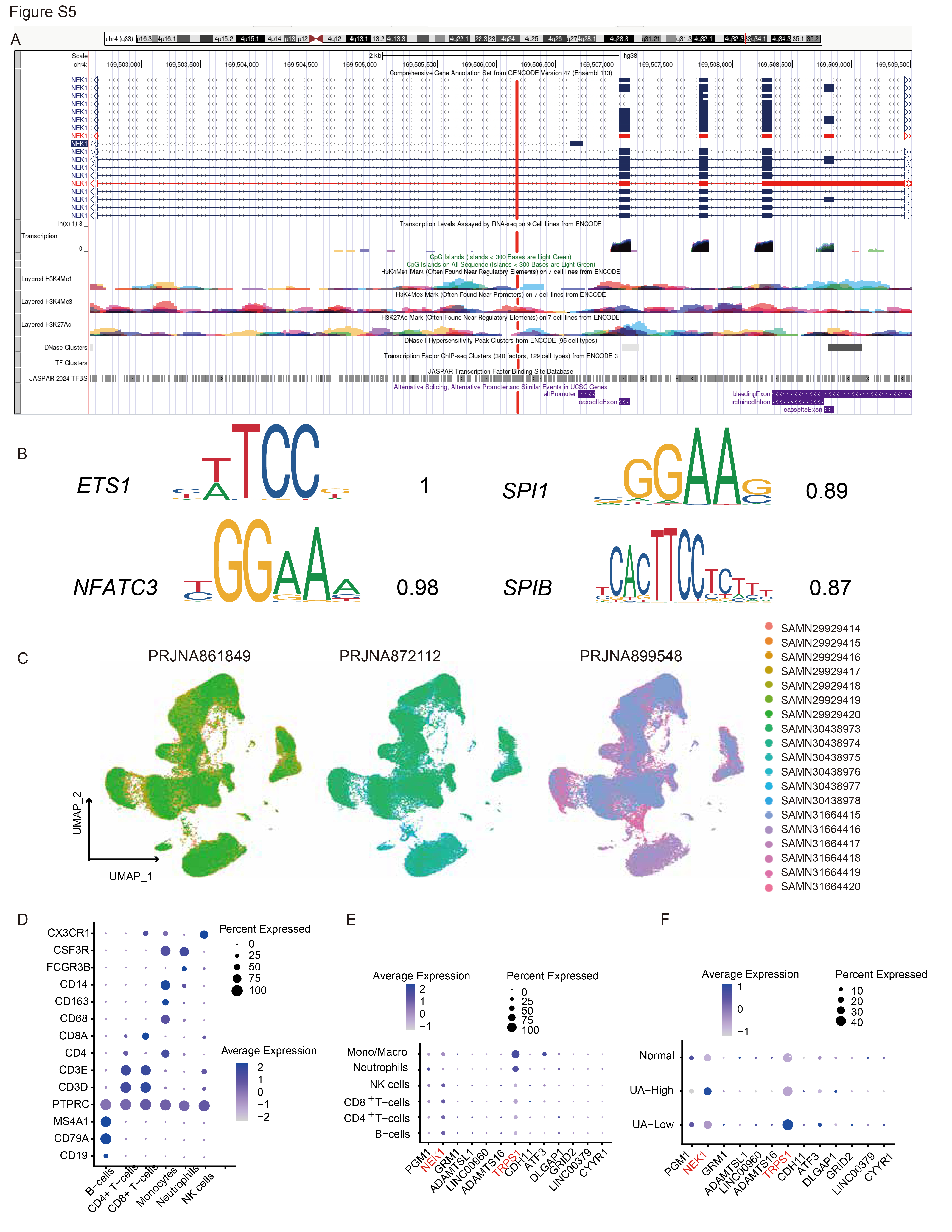
