## Supplementary material for "DNA methylation reveals molecular heterogeneity and urate excretion-associated subtypes of gout": Table S1

Table S1 Clinical characteristics of patients across different gout subtypes in the discovery cohort

| Parameters | RCB | RUE | ROL | RUN |
| --- | --- | --- | --- | --- |
| Number | 29 | 25 | 27 | 27 |
| Age, years | 38.00  (27.00-46.00) ^****^ | 37.00  (33.00-52.00) | 48.00  (42.50-54.00) | 55.00  (48.50-61.50) ^****^ |
| BMI, kg/m2 | 28.08  (25.69-29.88) | 27.45  (25.65-28.73) | 27.72  (24.98-30.24) | 25.51  (24.64-26.99) ^**^ |
| SBP, mmHg | 135.00  (124.00-142.00) | 128.00  (119.00-144.00) | 137.00  (128.50-144.50) | 129.00  (122.50-138.00) |
| DBP, mmHg | 85.00  (76.00-89.00) | 82.00  (78.00-89.00) | 88.00  (80.50-94.50) ^*^ | 84.00  (77.00-90.00) |
| SU, μmol/L | 583.00  (544.00-633.00) ^****^ | 536.00  (492.00-602.00) | 481.00  (432.00-533.50) ^*^ | 456.00  (424.00-512.00) ^***^ |
| Glu, mmol/L | 5.40  (5.00-5.86) | 5.31  (5.03-5.63) ^*^ | 5.62  (5.34-6.58) | 5.86  (5.32-6.23) |
| TG,  mmol/L | 1.86(1.31-2.54) | 1.85(1.24-2.59) | 1.51(1.14-2.02) | 1.60(1.33-2.29) |
| TC,  mmol/L | 4.90(4.64-5.43) | 4.75(4.48-5.40) | 5.16(4.32-6.14) | 5.05(4.67-5.54) |
| LDL-C,  mmol/L | 3.42(3.04-3.83) | 3.64(3.04-4.11) | 3.58(2.86-4.52) | 3.59(3.25-4.33) |
| HDL-C,  mmol/L | 0.98(0.84-1.12) | 1.06(0.95-1.19) | 1.15(1.00-1.44) | 1.14(1.07-1.20) |
| BUN, mmol/L | 4.50(3.60-5.00) | 4.00(3.60-4.50) ^*^ | 4.80(4.20-6.20) ^*^ | 5.00(4.00-6.00) |
| sCr, μmol/L | 134.00  (118.00-163.00) ^***^ | 122.00  (99.00-149.00) | 122.00  (102.00-143.50) | 89.00  (71.50-98.50) ^****^ |
| eGFR,  ml/min/1.73 m^2^ | 101.64  (90.20-117.44) ^**^ | 99.23  (84.12-116.09) | 93.68  (83.43-105.03) | 78.67  (68.50-84.45) ^****^ |
| Tophi, n (%) | 6 (21.43%),1 | 7 (28.00%) | 7 (26.92%),1 | 7 (25.93%) |
| Nephrolithiasis, n (%) | 4 (13.79%) | 4 (16.00%) | 4 (15.38%),1 | 5 (18.52%) |
| Hypertension, n (%) | 4 (13.79%) | 5 (20.00%) | 12 (46.15%) ^*^,1 | 10 (37.04%) |
| Cardiovascular disease, n (%) | 0 (0.00%) | 0 (0.00%) | 0 (0.00%),1 | 3 (11.11%) ^*^ |
| Diabetes mellitus, n (%) | 0 (0.00%) | 0 (0.00%) | 2 (7.69%),1 | 3 (11.11%) |
| Smoking, n (%) | 16 (55.17%) ^*^ | 6 (24.00%) | 10 (40.00%),2 | 5 (18.51%) |
| Drinking, n (%) | 20 (68.97%) | 20 (80.00%) | 21 (80.77%),1 | 19 (70.37) |
| Family history^a^, n (%) | 4 (13.79%) | 6 (24.00%) | 4 (14.81%) | 4 (14.81%) |
| FE_UA_, % | 4.45  (3.74-5.02) ^****^ | 3.58  (3.25-3.86) ^****^ | 6.33  (5.95-7.05) ^****^ | 6.04  (5.69-6.29) ^****^ |
| UUE, mg/d^-1^·(1.73 m^2^)^-1^ | 748.56  (745.17-758.95) ^**^ | 433.78  (430.40-434.83) ^****^ | 834.73  (793.76-844.04) ^****^ | 468.15  (451.33-498.25) ^***^ |

**Note:** Characteristics are presented as median (interquartile range). ^*^*P* ≤ 0.05; ^**^*P* ≤ 0.01; ^***^*P* ≤ 0.001; ^****^*P* ≤ 0.0001. ^a^ Family history of gout, hyperuricemia, diabetes, hypertension, or cardiovascular diseases. Categorical variables were analyzed using the *Fisher’s* exact test, while numerical variables were compared using analysis of the *Wilcoxon* test. A *P* value < 0.05 was considered statistically significant for all analyses. RCB, renal urate combined; RUE, renal urate underexcretion; ROL, renal urate overload; RUN, renal urate normal; BMI, body mass index; SBP, systolic blood pressure; DBP, diastolic blood pressure; SU, serum urate; Glu, fasting blood glucose; TG, triglyceride; TC, total cholesterol; LDL-C, low-density lipoprotein cholesterol; HDL-C, high-density lipoprotein cholesterol; BUN, blood urea nitrogen; sCr, serum creatinine; eGFR, estimated glomerular filtration rate; FE_UA_, fractional excretion of urinary urate; UUE, 24-h urinary urate amount.
