## Supplementary material for "DNA methylation reveals molecular heterogeneity and urate excretion-associated subtypes of gout": Table S2

Table S2 Clinical characteristics of patients across different gout subtypes in the validation cohort

| Parameters | RCB | RUE | ROL | RUN |
| --- | --- | --- | --- | --- |
| Number | 25 | 25 | 25 | 25 |
| Age, years | 38  (32.00-44.00) **^**^** | 39  (31.00-48.00) **^*^** | 42  (38.00-55.00) | 61  (54.00-66.00) **^****^** |
| BMI, kg/m2 | 27.93  (26.57-29.41) | 27.46  (24.49-29.41) | 27.68  (25.88-30.42) | 26.57  (24.62-28.04) **^*^** |
| SBP, mmHg | 137.00  (125.00-140.00) | 133.00  (122.00-147.00) | 146.00  (133.00-156.00) | 143.00  (128.00-154.00) |
| DBP, mmHg | 82.00  (78.00-92.00) | 87.00  (77.00-97.00) | 89.00  (80.00-102.00) | 87.00  (77.00-97.00) |
| SU, μmol/L | 600.00  (525.00-652.00) **^***^** | 532.00  (478.00-559.00) | 517.00  (463.00-584.00) | 497.00  (449.00-550.00) **^*^** |
| Glu, mmol/L | 5.65  (5.22-5.90) | 5.29  (5.15-5.67) **^**^** | 5.91  (5.38-7.09) **^**^** | 5.75  (5.27-6.27) |
| TG,  mmol/L | 1.95  (1.38-2.40) | 1.93  (1.41-2.58) | 2.10  (1.12-3.26) | 1.45  (1.18-2.12) |
| TC,  mmol/L | 4.88  (4.20-5.63) | 5.36  (4.43-5.76) | 5.30  (4.70-6.20) | 5.12  (4.55-5.76) |
| LDL-C,  mmol/L | 3.47  (2.78-3.79) | 3.50  (2.99-4.02) | 3.54  (3.26-3.99) | 3.62  (3.14-4.17) |
| HDL-C,  mmol/L | 1.02  (0.93-1.16) ^*^ | 1.12  (1.00-1.24) | 1.12  (0.95-1.29) | 1.13  (1.00-1.39) |
| BUN, mmol/L | 5.00  (4.20-5.70) | 4.10  (3.50-5.30) ^***^ | 5.50  (4.50-6.40) | 6.30  (4.80-7.00) ^**^ |
| sCr, μmol/L | 131.00  (101.00-159.00) ^**^ | 117.00  (96.00-148.00) | 117.00  (80.00-134.00) | 73.00  (64.00-86.00) ^****^ |
| eGFR,  ml/min/1.73 m^2^ | 95.13  (84.48-110.91) ^**^ | 87.99  (79.05-98.22) | 92.52  (69.16-98.92) | 72.26  (63.47-87.10) ^***^ |
| Tophi, n (%) | 8(32.00%) | 5(20.00%) | 8(32.00%) | 11(44.00%) |
| Nephrolithiasis, n (%) | 2(8.34%),1 | 0(0.00%) | 4(16.00%) | 4(16.00%) |
| Hypertension, n (%) | 10(41.67%),1 | 11(44.00%) | 18(72.00%) | 15(60.00%) |
| Cardiovascular disease, n (%) | 0(0.00%),1 | 0(0.00%) | 1(4.00%) | 2(8.00%) |
| Diabetes mellitus, n (%) | 1(4.17%),1 | 0(0.00%) | 4(14.00%) | 2(8.00%) |
| Smoking, n (%) | 12(48.00%),1 | 10(40.00%) | 16(64.00%) | 15(60.00%) |
| Drinking, n (%) | 13(68.42%),6 | 21(91.30%),2 | 19(79.17%),1 | 21(91.30%),2 |
| Family history^a^, n (%) | 20(80.00%) | 17 (68.00%) | 18(72.00%) | 19(76.00%) |
| FE_UA_, % | 4.42  (4.03-4.84) ^****^ | 4.12  (3.61-4.70) ^****^ | 5.95  (5.61-6.53) ^****^ | 5.86  (5.57-6.25) ^****^ |
| UUE, mg/d^-1^·(1.73 m^2^)^-1^ | 686.74  (652.32-757.58) ^***^ | 477.28  (396.63-528.52) ^****^ | 796.97  (751.92-972.81) ^****^ | 473.20  (414.66-567.62) ^****^ |

**Note:** Characteristics are presented as median (interquartile range). ^*^*P* ≤ 0.05; ^**^*P* ≤ 0.01; ^***^*P* ≤ 0.001; ^****^*P* ≤ 0.0001. ^a^ Family history of gout, hyperuricemia, diabetes, hypertension, or cardiovascular diseases. Categorical variables were analyzed using the *Fisher’s* exact test, while numerical variables were compared using analysis of the *Wilcoxon* test. A *P* value < 0.05 was considered statistically significant for all analyses. RCB, renal urate combined; RUE, renal urate underexcretion; ROL, renal urate overload; RUN, renal urate normal; BMI, body mass index; SBP, systolic blood pressure; DBP, diastolic blood pressure; SU, serum urate; Glu, fasting blood glucose; TG, triglyceride; TC, total cholesterol; LDL-C, low-density lipoprotein cholesterol; HDL-C, high-density lipoprotein cholesterol; BUN, blood urea nitrogen; sCr, serum creatinine; eGFR, estimated glomerular filtration rate; FE_UA_, fractional excretion of urinary urate; UUE, 24-h urinary urate amount.
