## Supplementary material for "DNA methylation reveals molecular heterogeneity and urate excretion-associated subtypes of gout": Table S3

| Index ID | BioProject | SRA BioSample | PMID | Tissue | Disease state | Uric acid (μmol/L) | Group |
| --- | --- | --- | --- | --- | --- | --- | --- |
| 1 | PRJNA861849 | SAMN29929419 | 36550178 | PBMCs | gout | 261.7 | UA-Low |
| 2 | PRJNA861849 | SAMN29929414 | 36550178 | PBMCs | gout | 285.5 | UA-Low |
| 3 | PRJNA861849 | SAMN29929417 | 36550178 | PBMCs | gout | 315.2 | UA-Low |
| 4 | PRJNA872112 | SAMN30438977 | 38063198 | PBMCs | acute gout | 335 | UA-Low |
| 5 | PRJNA861849 | SAMN29929418 | 36550178 | PBMCs | gout | 356.9 | UA-Low |
| 6 | PRJNA861849 | SAMN29929420 | 36550178 | PBMCs | gout | 422.3 | UA-Low |
| 7 | PRJNA899548 | SAMN31664418 | 37756161 | PBMCs | remission | 469.5 | UA-Low |
| 8 | PRJNA872112 | SAMN30438975 | 38063198 | PBMCs | acute gout | 486 | UA-Low |
| 9 | PRJNA872112 | SAMN30438975 | 38063198 | PBMCs | remission gout | 512 | UA-High |
| 10 | PRJNA872112 | SAMN30438974 | 38063198 | PBMCs | remission gout | 522 | UA-High |
| 11 | PRJNA899548 | SAMN31664416 | 37756161 | PBMCs | advanced remission | 522 | UA-High |
| 12 | PRJNA861849 | SAMN29929415 | 36550178 | PBMCs | gout | 523.4 | UA-High |
| 13 | PRJNA861849 | SAMN29929416 | 36550178 | PBMCs | gout | 523.4 | UA-High |
| 14 | PRJNA872112 | SAMN30438976 | 38063198 | PBMCs | acute gout | 533 | UA-High |
| 15 | PRJNA899548 | SAMN31664419 | 37756161 | PBMCs | remission | 613 | UA-High |
| 16 | PRJNA899548 | SAMN31664415 | 37756161 | PBMCs | advanced remission | 628 | UA-High |
| 17 | PRJNA899548 | SAMN31664420 | 37756161 | PBMCs | remission | 646 | UA-High |
| 18 | PRJNA872112 | SAMN30438973 | 38063198 | PBMCs | remission gout | 646 | UA-High |
| 19 | PRJNA899548 | SAMN31664417 | 37756161 | PBMCs | advanced remission | 676 | UA-High |

Table S3 Sample information and grouping of integrated single-cell RNA sequencing datasets

**Note:** PBMCs, Peripheral blood mononuclear cells.
