## Supplementary material for "DNA methylation reveals molecular heterogeneity and urate excretion-associated subtypes of gout": Table S4

| Index ID | Sample ID ^a^ | B cells | CD4^+^ T | CD8^+^ T | Mono-Macro | Neutrophils | NK |
| --- | --- | --- | --- | --- | --- | --- | --- |
| 1 | SAMN29929419 | 1042 | 4449 | 2508 | 2356 | 2 | 1647 |
| 2 | SAMN29929414 | 741 | 4380 | 1450 | 2427 | 2 | 809 |
| 3 | SAMN29929417 | 929 | 2563 | 1664 | 2104 | 12 | 1394 |
| 4 | SAMN30438977 | 267 | 594 | 466 | 2801 | 23 | 615 |
| 5 | SAMN29929418 | 660 | 4430 | 1293 | 1887 | 2 | 1450 |
| 6 | SAMN29929420 | 618 | 4408 | 1239 | 2652 | 43 | 1538 |
| 7 | SAMN31664418 | 921 | 7012 | 2915 | 208 | 0 | 2716 |
| 8 | SAMN30438975 | 211 | 838 | 846 | 1662 | 91 | 502 |
| 9 | SAMN30438975 | 297 | 1821 | 1100 | 1673 | 12 | 642 |
| 10 | SAMN30438974 | 298 | 1337 | 866 | 1453 | 10 | 674 |
| 11 | SAMN31664416 | 298 | 1337 | 866 | 1453 | 10 | 674 |
| 12 | SAMN29929415 | 935 | 3475 | 1886 | 560 | 0 | 1708 |
| 13 | SAMN29929416 | 620 | 4540 | 1135 | 3063 | 5 | 1610 |
| 14 | SAMN30438976 | 783 | 8650 | 2154 | 261 | 0 | 1437 |
| 15 | SAMN31664419 | 315 | 6712 | 2339 | 2092 | 1 | 1622 |
| 16 | SAMN31664415 | 746 | 4871 | 3007 | 193 | 0 | 3443 |
| 17 | SAMN31664420 | 635 | 3849 | 2688 | 324 | 5 | 2453 |
| 18 | SAMN30438973 | 921 | 7012 | 2915 | 208 | 0 | 2716 |
| 19 | SAMN31664417 | 297 | 1821 | 1100 | 1671 | 12 | 642 |

### Table S4 Numbers of cells per sample after quality control

**Note：** Mono-Macro, Monocytes and Macrophages; NK, Nature Kill cells. ^a^ Data were obtained from GSE211783 (NCBI GEO Database), GSE217561 (NCBI GEO Database), and PRJNA861849 (NCBI SRA Database)
