## Supplementary material for "DNA methylation reveals molecular heterogeneity and urate excretion-associated subtypes of gout": Table S9

Table S9. Genomic locations and functional features of 18 methylation loci in the gout model.

| **Locus** | **Gene** | **Gene type** | **Genomic location** | **Associated functions** | **Phenotypes** |
| --- | --- | --- | --- | --- | --- |
| chr1.63649272 | *PGM1* | Protein Coding | Promoter (2-3kb) | This enzyme participates in both the breakdown and synthesis of glucose (PubMed:17924679, 25288802) | Bone tissue density, brain attribute, phosphoglucomutase-1 measurement |
| chr10.110935823 | *RPL13AP6* | Pseudogene | Promoter (1-2kb) | Ribosomal Protein L13a Pseudogene 6. No data available | Sarcosine measurement |
| chr4.169506188 | *NEK1* | Protein Coding | Promoter (<=1kb) | In response to injury that includes DNA damage, NEK1 phosphorylates VDAC1 to limit mitochondrial cell death (PubMed:20230784). Involved in cilium assembly (PubMed:21211617). | Schizophrenia, educational attainment, smoking initiation |
| chr6.146231911 | *GRM1* | Protein Coding | Intron | May participate in the central action of glutamate in the CNS, such as long-term potentiation in the hippocampus and long-term depression in the cerebellum (PubMed:24603153, 28886343, 7476890). | body height, lobe attachment, appendicular lean mass |
| chr9.18214919 | *ADAMTSL1* | Protein Coding | Intron | This gene encodes a secreted protein and member of the ADAMTS (a disintegrin and metalloproteinase with thrombospondin motif) family. | bone tissue density, body height, abnormality of refraction |
| chr17.53584328 | *KIF2B* | Protein Coding | Intron | Has microtubule depolymerization activity (PubMed:17538014). Plays a role in chromosome congression (PubMed:23891108). | smoking initiation, educational attainment, body height |
| chr3.75668878 | *LINC00960* | lncRNA | Intron | No data available. | alzheimer disease, family history of alzheimer’s disease |
| chr16.63463024 | *LOC105371308* | ncRNA | Intron | No data available. | vascular disease,educational attainment,neuritic plaque measurement |
| chr5.5203251 | *ADAMTS16* | Protein Coding | Intron | No data available. | bone tissue density, level of a disintegrin and metalloproteinase with thrombospondin motifs 16 in blood, abnormal result of function studies |
| chr8.115804703 | *TRPS1* | Protein Coding | Intron | Transcriptional repressor. Executes multiple functions in proliferating chondrocytes, expanding the region of distal chondrocytes, activating proliferation in columnar cells and supporting the differentiation of columnar into hypertrophic chondrocytes. | bone tissue density, body height, sialic acid-binding ig-like lectin 8 measurement |
| chr16.64957802 | *CDH11* | Protein Coding | Intron | Required for proper focal adhesion assembly (PubMed:33811546). Involved in the regulation of cell migration (PubMed:33811546). | blood protein amount, body height, protein measurement |
| chr1.212592288 | *ATF3* | Protein Coding | Intron | This protein binds the cAMP response element (CRE) (consensus: 5'-GTGACGT[AC][AG]-3'), a sequence present in many viral and cellular promoters. | cleft lip, cleft palate, leukocyte quantity |
| chr18.4444736 | *DLGAP1* | Protein Coding | Intron | Part of the postsynaptic scaffold in neuronal cells. | body height, age at menarche, testosterone measurement |
| chr4.92693189 | *GRID2* | Protein Coding | Intron | Cation-selective ion channel activity can be triggered by GRM1 in Purkinje cells (PubMed:24357660, 27276689). | bone tissue density, level of coiled-coil domain-containing glutamate-rich protein 2 (human) in blood, body height |
| chr13.91154417 | *LINC00379* | lncRNA | Distal Intergenic | No data available. | body height, glypican-5 measurement, anthropometric measurement |
| chr2.89792905 | *KMT2CP4* | Pseudogene | Distal Intergenic | No data available. | calcium measurement, complement factor d measurement |
| chr21.26647755 | *CYYR1* | Protein Coding | Distal Intergenic | No data available. | viral load, body height, strand of hair color |
| chr3.96009846 | *MTHFD2P1* | Pseudogene | Distal Intergenic | No data available. | bone tissue density,body heightm,enteritis |
