## Supplementary material for "DNA methylation reveals molecular heterogeneity and urate excretion-associated subtypes of gout": Table S10

Table S10 Predicted Motifs potentially associated with the methylation site Chr4:169506188

| Index ID | Matrix ID | Name | Score | Relative score | Position | Strand | Predicted sequence |
| --- | --- | --- | --- | --- | --- | --- | --- |
| 1 | MA0098.1 | MA0098.1.ETS1 | 7.797 | 1 | chr4:169506188 | - | TTTCCG |
| 2 | MA0625.3 | MA0625.3.NFATC3 | 9.505 | 0.984 | chr4:169506188 | + | CGGAAA |
| 3 | MA0080.1 | MA0080.1.SPI1 | 5.834 | 0.879 | chr4:169506188 | + | CGGAAA |
| 4 | MA0081.3 | MA0081.3.SPIB | 12.925 | 0.870 | chr4:169506188 | - | GCATTTCCCCTTT |
| 5 | MA1940.2 | MA1940.2.ETV2::DRGX | 10.370 | 0.860 | chr4:169506188 | + | CGGAAAGAATTC |
| 6 | MA1117.1 | MA1117.1.RELB | 8.583 | 0.848 | chr4:169506188 | - | GCATTTCCCCT |
| 7 | MA1944.2 | MA1944.2.ETV5::DRGX | 9.089 | 0.827 | chr4:169506188 | + | CGGAAAGAATTC |
| 8 | MA1121.2 | MA1121.2.TEAD2 | 6.171 | 0.822 | chr4:169506188 | - | GCATTTC |
| 9 | MA0673.2 | MA0673.2.NKX2-8 | 1.701 | 0.805 | chr4:169506188 | - | GCATTTCC |
| 10 | MA0076.2 | MA0076.2.ELK4 | 3.949 | 0.802 | chr4:169506188 | - | GCATTTCCCCT |
| 11 | MA0098.1 | MA0098.1.ETS1 | 7.797 | 1 | chr4:169506188 | - | TTTCCG |
